# Allele-specific differential regulation of monoallelically expressed autosomal genes in the cardiac lineage

**DOI:** 10.1101/2022.02.01.478691

**Authors:** Gayan I. Balasooriya, David L. Spector

## Abstract

Each mammalian autosomal gene is represented by two alleles in diploid cells. Yet, no insights have been made in regard to allele-specific regulatory mechanisms of autosomes. Here we used allele-specific single cell transcriptomic analysis to elucidate the establishment of monoallelic gene expression in the cardiac lineage. We found that monoallelically expressed autosomal genes in mESCs and mouse blastocyst cells are differentially regulated based on the genetic background of the parental alleles. However, the genetic background of the allele does not affect the establishment of monoallelic genes in differentiated cardiomyocytes. Additionally, we observed epigenetic differences between deterministic and random autosomal monoallelic genes. Moreover, we also found a greater contribution of the maternal versus paternal allele to the development and homeostasis of cardiac tissue and in cardiac health, highlighting the importance of maternal influence in male cardiac tissue homeostasis. Our findings emphasize the significance of allele-specific insights into gene regulation in development, homeostasis and disease.

## Introduction

The diploid eucaryotic genome requires precise spatial-temporal regulation during development and homeostasis. Autosomal gene transcription can be carried out in three states; bi- allelic, deterministic monoallelic (DeMA – the allele selection is predetermined. e.g. imprinted genes), and random autosomal monoallelic (RaMA – the allele selection is not predetermined. e.g. each allele has an equal chance to become the active or the inactive allele). Alteration to the homeostatic expression from the selected DeMA or RaMA gene may result in allele heterogeneity, thus may partly explain the variable penetrance of disease-associated mutations caused by monoallelic genes. Hence, extensive investigation into the establishment and mechanism of monoallelic gene expression can provide insights into development, homeostasis, and disease. RaMA gene expression is widely established in early mammalian development^1–5^. This gene expression paradigm has been explored using bulk cell transcriptomic data generated from *in vitro* F1 mouse embryonic stem cell (mESC) derived neural progenitor cells^2, 3^, however, it has not been extensively investigated in cells or tissues committed to different lineage differentiation paradigms at single cell resolution. Sex-biased autosomal gene expression has been recently reported, but allele-exclusive expression has not yet received much attention^6, 7^. The regulatory bias based on the alleles’ origin and alleles’ genetic background in autosomal monoallelic genes has been overlooked, as dogma implies that there is no distinction between maternal and paternal alleles based on the similarity of the nucleic acid chemistry. Thus, here we sought to investigate the allele-specific expression profiles of monoallelic genes in cells of the cardiac lineage.

## Results

### Allele-bias of DeMA gene expression

The establishment of monoallelically expressed genes in cardiac lineage cell types is currently unknown. We initially implemented an allele-specific bulk-RNA-seq approach on highly morphogenic (C57Bl/6J-maternal x CAST/EiJ-paternal) male F1 mouse embryonic stem cells (mESC) derived from three blastocysts and *in vitro* derived heterogenous cardiac precursor cell cultures (CPCCs) from those three F1 mESC clones. CPCCs are predominantly enriched with cardiac stem cells (CSCs) and cardiac progenitor cells (CPCs) (Fig.1a & Extended Data Fig.1a-c), but also contain an additional three cell types, including *Mef2c* positive cardiac precursor cells (Fig.2c & Extended Data Fig.2b). Since the CPCCs contain lineage committed naive and fully differentiated CSCs and CPCs, these cultures exhibit a transcriptional profile similar to mESCs and a transcriptional profile specific to cardiac lineage cells (Fig.1b & Extended Data Fig.1c, **d).** To examine the parental origin of the transcripts, we systematically implemented a rigorous allele-specific gene filtering algorithm^8^ so that the significantly expressed autosomal genes from each parental background could be identified as deterministic maternal (mat-mono), deterministic paternal (pat-mono) and bi-allelically expressed gene cohorts (Fig.1c, d, Extended Data Fig.1 **e).** In this rigorous filtering approach, we only considered a gene to be a DeMA gene if it is solely expressed from one parental background in all three clonal biological replicates. The intersect of all mESC and CPCCs mat-mono and pat-mono genes showed no allele switching of expressed genes between the cell types (except for three outliers) (Fig.1e). Interestingly, during mESC to CPCC differentiation, we observed that the 994 monoallelically expressed genes in mESCs had changed their expression to biallelic status and the 468 biallelic genes in ESCs had changed to monoallelic status (Extended Data Fig. 1f, **g).** The newly identified 266 genes out of 1496 monoallelic genes in CPCCs function in biological processes relevant to cardiac lineage cells (Extended Data Fig. 1h) indicating the biological relevance in the establishment of DeMA genes in cardiac lineage cells.

**Fig. 1.**
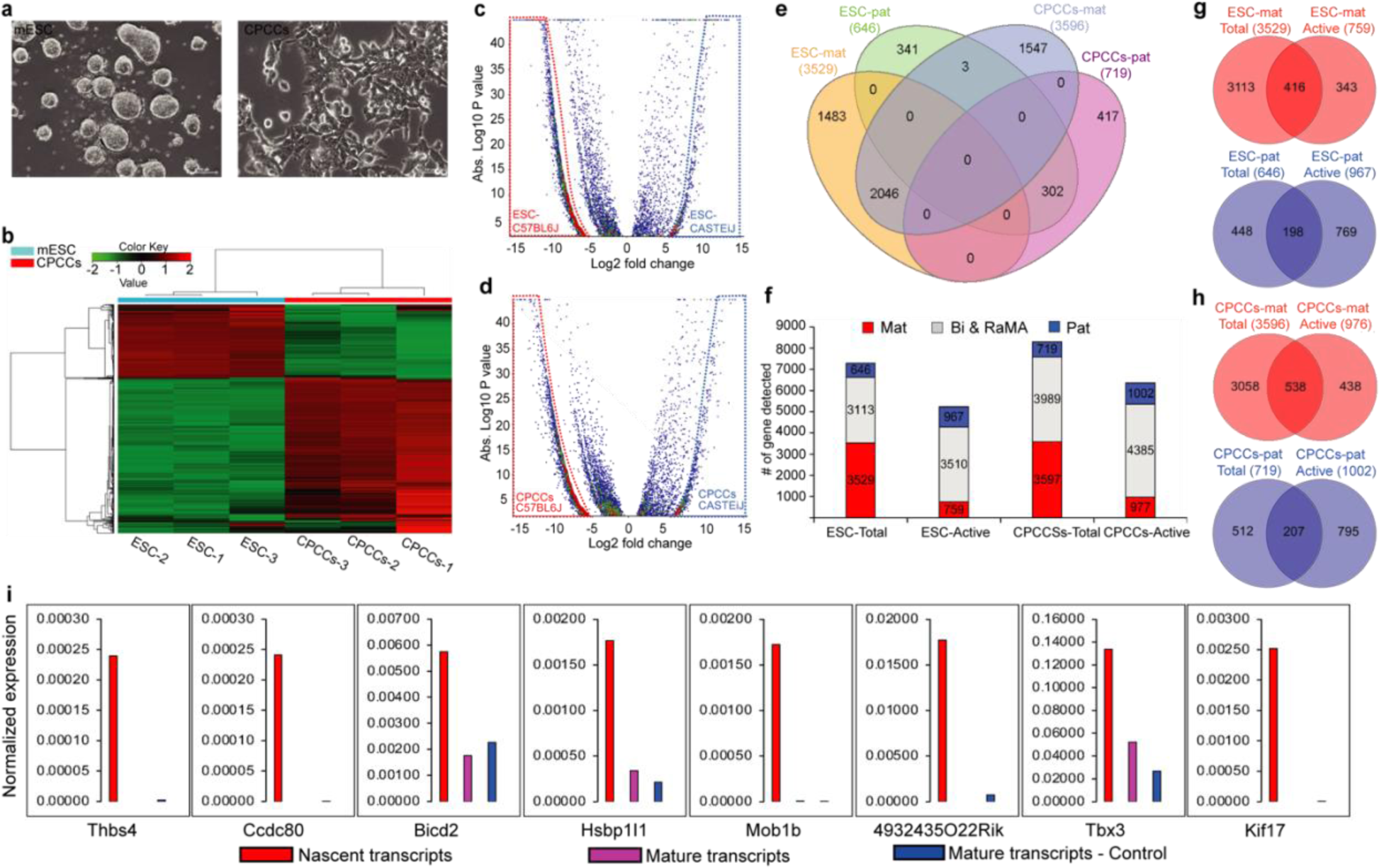
Allelic expression of autosomal monoallelic genes. **a**, F1 mESCs grown in 2i medium and in vitro derived CPCCs cultures. **b,** Hierarchical cluster heat map of F1 mESC and CPCCs. Gene wise dispersion by conditional maximum likelihood (Log2(CPM)) was assessed using EdgeR pipeline setting the cut-off z-score at four. Bulk-RNA-Seq performed with biological triplicates. **c,d**, Volcano-plot showing maternally and paternally expressed genes in mESCs and CPCCs. Log2 fold changes of the (-)X axis coordinates are deterministic mat-mono genes (red dotted line area) and the (+)X axis coordinates are deterministic pat-mono genes (blue dotted line area). Differential gene expression analysis was performed using EdgeR pipeline. **e,** Intersects of mESC and CPCCs monoallelic genes. Allele-specific monoallelic genes: Some are common between mESCs and CPCCs, the remainder are unique to each cell type and do not switch alleles (except three outliers). f, Mat-mono and pat-mono establishment is imbalanced in both mESCs and CPCCs. The number of actively transcribing versus mature transcripts differ. g,h, Intersects of actively transcribing genes versus genes with mature transcripts. Higher proportion of genes contain either only the mature transcripts or only the nascent transcripts. i, qRT-PCR analysis for representative actively transcribing monoallelic genes lacking mature transcripts. Nascent and mature transcripts were assessed using a Click-iT™ assay and mature transcript–control was assessed on total RNA used in the Click-iT™ assay (n=3). Nascent transcripts expressed were normalized to nascent Actb transcripts expressed and the mature transcripts expressed were normalized to mature Actb transcripts expressed.

To investigate whether distinct maternal versus paternal DeMA gene expression is due to the parent-of-origin of the alleles or the alleles’ genetic background, we re-analyzed three publicly available RNA-seq data sets (two mESC bulk RNA-seq (GSM2405897)^9^, one bulk RNA-seq from 32-cell blastocysts (GSE152103)^10^ and one scRNA-seq data set from 32-cell blastocysts (GSE80810)^11^) generated from C57BL/6J x CAST/EiJ reciprocal crosses. Though the number of genes expressed in the same cell types in the reciprocal crosses is the same, we found that the divergence of the imbalance of DeMA gene expression from maternal and paternal alleles is due to the genetic background of the parental alleles. We found that the C57BL/6J allele is preferentially expressed in *in vitro* mESCs and uncultured cells from blastocysts, which to our knowledge, was not previously reported in any studies using crosses between different genetic backgrounds (Extended Data Fig.1i-m).

We observed that in mESCs and CPCCs, the number of mat-mono genes expressed was 5.5-fold and 5.0-fold higher than the number of pat-mono genes, respectively (Fig.1f). To test whether this imbalanced allelic expression can be explained by differential transcription bursting frequencies in each genetic background, we assessed the active transcription status of the maternal and paternal alleles in our RNA-seq data *in silico*. We observed in both mESCs and CPCCs that the number of actively transcribed mat-mono genes is lower than the total mat-mono gene number (Fig.1f) and that no nascent transcripts were detected for a subset of mature mat-mono mRNAs (ESCs: 88.2% and CPCCs: 85.7% from the total RNAs) (Fig.1g, h), perhaps due to those genes’ longer time-lag between transcriptional bursts. Conversely, a significant number of pat-mono actively transcribing genes were lacking, or exhibited a significantly reduced number of, mature transcripts (ESCs: 79.5% and CPCCs:79.3% from the total active RNAs) (Fig.1g, h). To validate this unexpected observation, we performed nascent and mature RNA specific qRT-PCR for several candidate genes (Fig.1i). qRT-PCR results were in accordance with our *in-silico* observation and thus we suggest this is possibly due to delayed pre-mRNA processing or rapid mRNA decay^12, 13^, or a yet unknown nascent mRNA perpetuation mechanism of those mRNA species.

### Allele-biased DeMA gene expression in cardiac lineage commitment

To investigate allele-biased DeMA gene expression further in lineage committed and terminally differentiated somatic cardiac cells, we conducted allele-specific scRNA-seq analysis in mESCs (215-cells) and cells from *in vitro* derived CPCCs (175-cells) and cardiac bodies (CBs, 232-cells) (Extended Data Fig.1a & Video-1). Pseudo-time trajectory analysis confirmed the linear-lineage correlation of the mESCs, CPCCs and CBs (Fig.2a). Next, the cellular heterogeneity of each point was defined by a systematic cell-cluster analysis approach (Fig.2b-d, Extended Data Fig.2a-c). From those cell-clusters we investigated allele-specific expression in seven known cell types: G1/S-mESCs, CSCs, CPCs, CF-cardiac fibroblasts, CM-cardiomyocytes, EC- endocardium cells, and PE-pericardial cells. Cardiac lineage cell-clusters did not further sub-cluster in cell cycle analysis. In agreement with the observation from bulk-RNA-seq data (Fig.1f,g), allele-specific scRNA-seq analysis confirmed that the distinctive DeMA gene expression pattern from parental genomes remains preserved between mESCs and six cardiac lineage cell types (Fig.2e, Extended Data Fig.2d). In addition, active mat-mono and pat-mono genes exhibit imbalanced allelic expression frequencies, either higher in pat-mono genes (e.g.CSC, CPC, EC and PE) or higher in mat-mono genes (e.g., ESCs, CF, CM) (Extended Data Fig.2e). Further, a higher proportion of mat-mono genes are shared between cardiac cell types than pat-mono genes and they rarely exhibit allele-switching (Fig.2f).

**Fig. 2.**
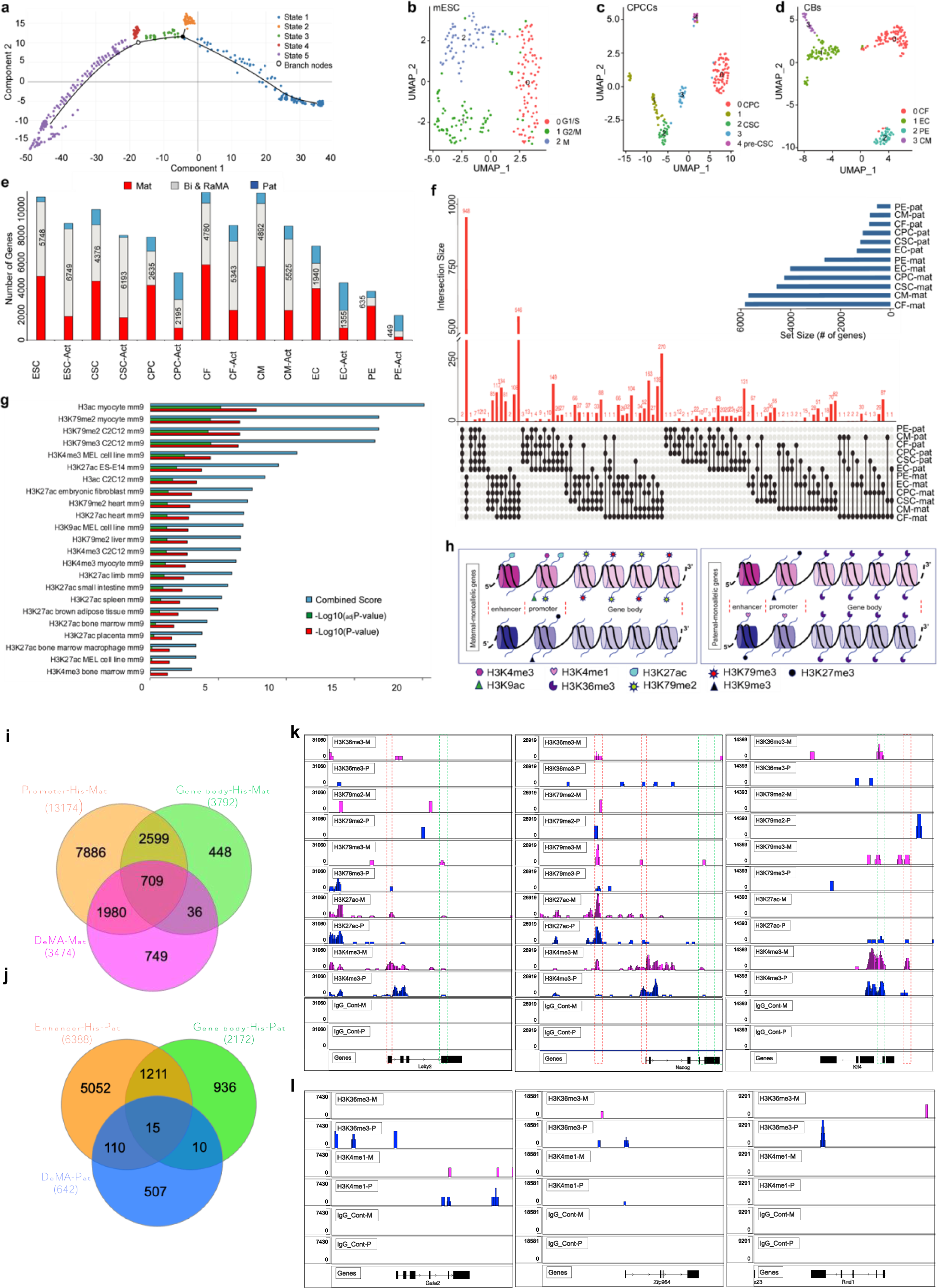
Allele expression in cardiac cell types and allele-specific differential epigenetic regulation of monoallelic genes. **a**, Pseudo time trajectory analysis for mESCs, CPCCs and CBs scRNA-Seq data. Trajectory shows the linear-lineage correlation of three experimental points. State 1: mainly mESCs, State 2,3 and 4: mainly CPCCs, State 5: mainly CBs. Nodes of the trajectory indicate the terminating points between the experimental points. **b**-**d**, Uniform Manifold Approximation and Projection (UMAP) based systematic cell cluster analysis for mESC, CPCCs and CBs scRNA-Seq data points. UMAP plots show three clusters for mESC, five for CPCCs (Cluster 0: CSC, Cluster 2: CPC) and four for CBs (Cluster 0: CF, Cluster 1: CM, Cluster 2: EC, Cluster 3: PE). **e**, Number of bi-allelic and DeMA genes expressed in seven cell types. For each cell type, the number of bi-allelic and DeMA genes expressed are shown for two transcript categories: mature transcripts and nascent transcripts. **f**, Intersects of all maternal and paternal monoallelic genes in cardiac lineage cell types. Significantly higher number of mat-mono genes are shared between all cardiac cell types than pat-mono genes. Allele switching less commonly occurred between maternal and paternal backgrounds; 100 intersects are shown. Nodes show the intersected gene size and the line shows the cell types that are sharing the intersected genes. **g**, Histone signatures of mat-mono genes. Histone marks enrichment represented as Combined score, -Log_10_(adjP-value), -Log_10_(P-value) (see methods) for each signature. **h**, Schematic representation of the extrapolation of the histone mark association for DeMA gene alleles. Maternal allele: Purple colored. Paternal alleles: Blue colored. **i**,**j**, Allele specific histone signatures in F1 mESCs from CUT&Tag assay compared with maternal and paternal DeMA genes. **i**, Histone signatures in promoters (H3Kme3, H3K27ac and H3K79me2) and gene bodies (H3K79me2, H3K79me3 and H3K36me3) of maternal allele comparison with maternal DeMA genes (DeMA-Mat) in RNA-seq data from bulk ESCs. From the total maternal DeMA genes, 78.4% of maternal DeMA genes are exclusively enriched in maternal allele histone signatures. **j**, As *in silico* predicted in ENCODE data (at the low probability p=0.03, compared to maternal genes), the paternal DeMA genes (DeMA-Pat) from bulk RNA-seq data from mESCs do not show higher correlation (21%) with enhancer (H3K4me1) or gene body (H3K36me3-form Fig.2h) association in CUT&Tag data. **k**, Illustration of five allele-specific histone associations in maternal DeMA genes in mESCs. *Nanog*, *Klf4* and *Lefty2*, known pluripotent associated genes. Red dotted lines indicate the enhancer/promoter regions and green dotted lines indicate the gene body region and histone association. For example, H3K4me3, H3K79me3 and H3K36me3 histone association in ∼ +5 kb region of Nanog TSS in maternal allele is shown (*Nanog* is maternaly expressed in our study model). **i**, In paternally expressed *Gata2*, *Zfp964* and *Rind1* genes, only the paternal alleles indicate the enhancer and gene body histone signatures.

To further validate our data sets, we investigated whether known monoallelic genes expressed in the heart were included in our CM specific DeMA gene set. We observed the allele-specific expression of several known imprinted genes including *H19*, *Snrpn*, *Cobl* and *Dlk1* agreeing with our data set, but some imprinted genes, such as *Peg10*^14^, was observed to be biallelically expressed in CMs. It has been shown that some of the imprinted monoallelic genes show clustered gene expression. Although it was previously demonstrated that *Ubea3a* exhibited clustered gene expression with *Snrpn*, we did not see *Ubea3a* expressed in our single-cell CMs specific gene set. Instead, we detected that *Snurf,* which is within the *Snrpn* cluster, is also maternally expressed. As previously reported, we also observed the maternally expressed *Begain*, which is positioned at the Chr12 centromeric boundary, physically distanced ∼383 kb to the *Dlk1* imprinted gene at the *Dlk1*-*Meg3* imprinted cluster^15–17^. However, we saw no *Meg3* expression in CMs perhaps due to the cell type specific gene expression of *Meg3* in the heart. Some of the differences we detect between our DeMA genes and others may be due to the different technical approaches we used (e.g. single cell vs bulk tissue).

### DeMA genes contain distinctive epigenetic marks

Allele-specific gene expression may be explained by the distinctive epigenetic status of the alleles^18, 19^. However, the histone signatures of cardiac monoallelic genes have been largely unexplored. Previous studies suggest that H3K27me3 and H3K36me3 are the primary histone signatures of monoallelic genes^20–22^, but the signatures of the active versus silent allele were not examined individually. To address the allele-specific histone signatures of DeMA genes, we compiled all the mat-mono and pat-mono gene candidates identified in the seven cell types with ENCODE histone modification data to probe their common histone signatures (Extended Data Table 1). This analysis revealed that the maternal and paternal monoallelic alleles are enriched for different and unique histone signatures (Fig.2g, Extended Data Fig.2f). H3ac, H3K79me2/3, H3K4me3, H3K27ac and H3K9ac are the signatures significantly enriched on mat-mono genes. H3K27ac marks the active gene enhancers^23, 24^. Bivalent enrichment of H3K4me3 and H3K9ac and; H3K27ac and H3K79me2 signatures occupy highly active gene promoters. In contrast, the same promoters contain H3K27me3 and H3K9me3 histone marks (lowly abundant in our data, Extended Data Table-1) when those same genes become transcriptionally silent in cardiomyocytes^25^, suggesting the possibility of H3K27me3 and H3K9me3 silent histone signatures marking the silent paternal alleles of maternally expressed genes. In our pat-mono genes, the most significant signature is H3K27me3, a silent gene histone signature at the promoter or downstream of the transcription start sites (TSSs). H3K4me1 (enhancer mark) is also present (less significance: combined score = 0.33) in our pat-mono genes, suggesting a model where H3K4me1 and H3K27me3 perhaps bivalently occupy the poised enhancers^26^ and H3K4me1, the unimodal histone mark, occupies the poised promoter of the active paternal allele. Heterochromatin associated H3K9me3 and H3K36me3 histone marks are lowly abundant in our pat-mono genes (Extended Data Table 1). We speculate that the repressed maternal alleles of pat- mono genes are either marked by H3K27me3 and H3K9me3 at the promoter regions simultaneously with H3K36me3 in the genes’ body, or the enhancer/promoters maintain their silent state, perhaps by another mechanism such as miRNA, siRNA or DNA secondary structures^27, 28^. We also observed that the average transcription levels were higher in mat-mono genes than in the pat-mono genes in mESCs, CFs and CMs (Extended Data Fig.2e), but in the CSCs, CPCs, ECs and PEs, the average transcription levels were higher in pat-mono genes than mat-mono genes. This may be the result of the enhancer/promoter association of different chromatin signatures in different cell types, which means for our data that in mESCs, CFs and CMs, the enhancer associated H3K27ac mark may promote a higher rate of mat-mono gene transcription, while in pat-mono genes H3K4me1 poised promoters may drive steady-state transcription, resulting in a lower transcription levels^29, 30^. In contrast, in CSCs, CPCs, ECs and PEs, unimodal H3K4me1 signatures on promoters (proximal to TSS) may result in higher pat- mono gene transcription, a muscle cell development gene specific epigenetic mechanism^31^. For the most part, allele specificity of monoallelic genes does not change between cardiac cell types although a few outliers exhibit a switch of the parental origin but remained monoallelic (Fig.2f). This tightly regulated mechanism, in part in mat-mono genes, may be driven by H3K79me2/3, which may permanently ‘bookmark’ active alleles through mitosis^32^ as the H3K79me2/3 mark occupies active gene bodies through mitosis but is not involved in gene activation^25, 33–35^. Taken together, incorporating our data with the published data, we propose an allele-specific histone signature model for DeMA genes as illustrated in Fig.2h.

To experimentally validate the histone signatures in our proposed model, as a proof of principle, we next assessed the six allele specific histone signatures in mESCs (Extended Data Fig.2g) using a CUT&Tag^36^ assay. As expected, we detected strong H3K4me3 and H3K27ac signals around the TSS of the genes (Extended Data Fig.2h). However, when we assessed the allele-specific enrichment (Extended Data Fig.2i, j), we noticed that two peaks were present upstream and downstream of the TSSs, probably indicating proximal enhancers/promoters at the 5’ of the TSSs and at the promoters 3’ of the TSSs. H3K4me1, the only enhancer mark on the paternal alleles in mESCs was noticeably enriched upstream of the TSSs as well as in the gene bodies, probably at intergenic enhancers. H3K79me2 and H3K79me3 known to mark gene bodies, was also observed at the TSSs, suggesting that those signatures may be involved in assisting the enhancer-promoter interactions of actively transcribing genes^37, 38^. Moreover, we found that H3K36me3, a signature of actively transcribing gene bodies, was enriched at the promoter regions (Extended Data Fig.2g), but was not strongly present at DeMA gene promoters, matching to the histone signatures we modeled for mESC (Extended Data Fig.2g) DeMA genes (Extended Data Fig.2i, j).

Next, to validate the histone signature model we proposed for DeMA genes, we assessed our mESCs’ maternal and paternal gene cohorts from bulk-RNA-seq data in allele-specific histone signatures from the CUT&Tag assay. We found 78.4% of our maternal DeMA genes fell within the maternal-specific histone signatures (Fig.2i & k). However, the less probabilistically significant (P=0.03) *in-silico* predicted paternal allele-specific histone signatures (Extended Data Fig.2f – lower panel) only matched with 21% of the genes we identified in our CUT&Tag assay (Fig.3j & l). We reason that this is a probabilistic issue of the analysis of paternal DeMA genes caused by the predicted low-number (n=2) of parental specific histone marks from *in-silico* analysis. Therefore, our histone signature model, at least in part, is supported by the histone signatures associated with the maternal DeMA genes.

**Fig. 3.**
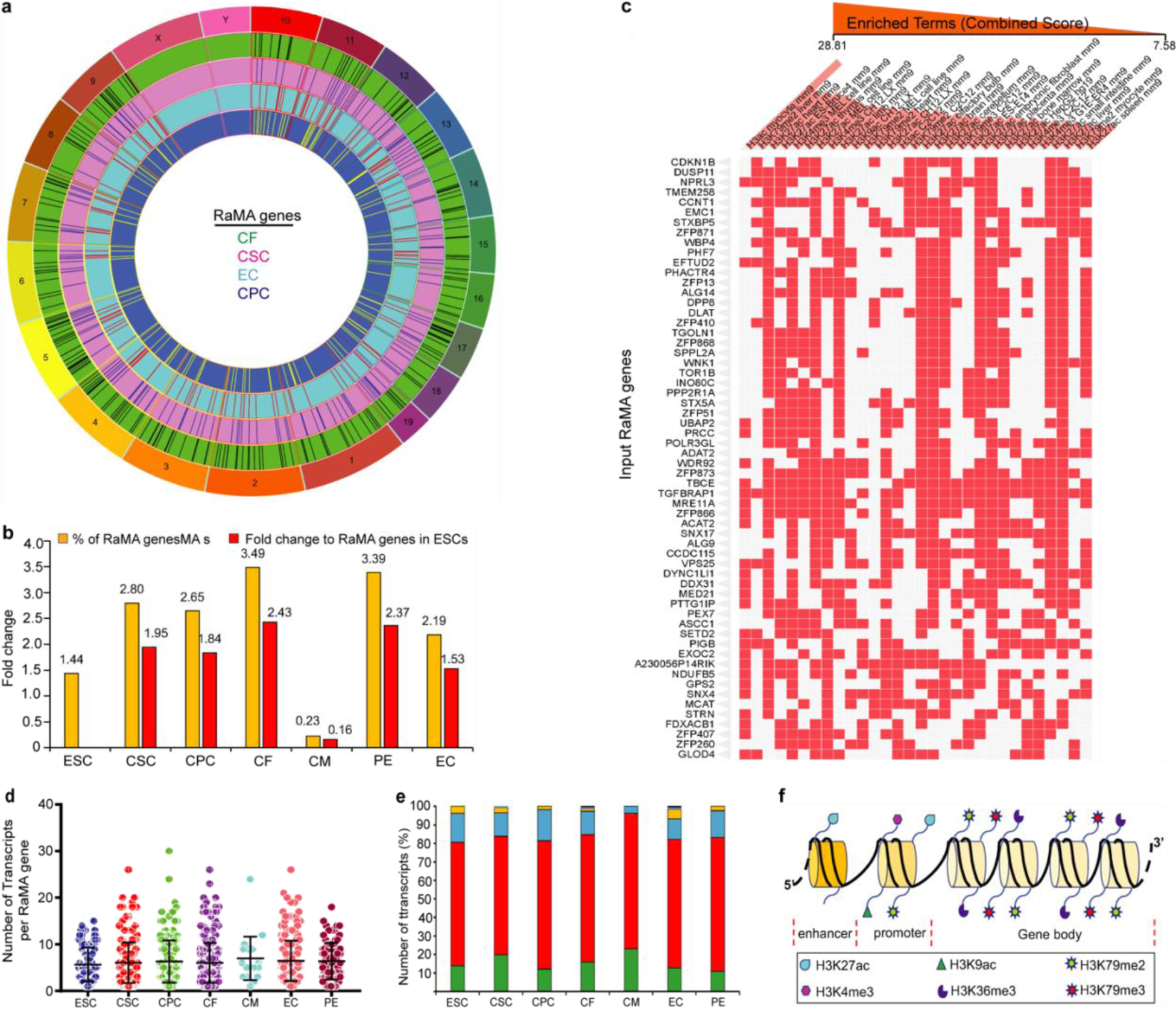
RaMA genes exhibit lineage specificity and expression is regulated by distinct histone modifications to monoallelic genes. **a**, Spatial arrangement of CSC, CPC, CF and EC RaMA genes in the autosomes. Columns in each circle represent the gene positions. **b**, Percentages and the fold changes of RaMA genes upon ESC differentiation (scRNA-seq). Percentages are calculated to non-monoallelic genes and the fold changes are with reference to RaMA gene number in mESCs. **c**, Histone signatures of RaMA genes from seven cell types. Thirty terms and 60 RaMA genes with the highest significance (Combined score) are shown. Enriched terms are columns (column order is the sum) and the input genes are the rows. **d**, Number of transcripts in RaMA genes in seven cell types. Unpaired t-test was performed and the error bars presents the standard deviations of the mean values. **e**, The number of known protein coding transcripts per gene in protein coding RaMA genes in seven cell types. Only experimentally validated proteins matched to merged Ensembl/Havana transcripts were considered as protein coding transcripts. **f**, Schematic representation of the extrapolation of the histone mark association for expressed RaMA gene alleles.

### Context dependent RaMA gene expression

RaMA genes stochastically select the expressed and repressed alleles. To assess whether RaMA genes share the same regulatory mechanism as DeMA genes, we first identified RaMA genes in seven cell types using a rigorous gene filtering method (Extended Data Fig.3a). In line with previous reports, we observed that RaMA genes are not clustered, rather they are randomly positioned in the genome^2–4, 39^ (Fig.3a). An increased number of RaMA genes has been reported in *in vitro* cell differentiation paradigms^2, 3^. To determine whether an increase in RaMA expression occurs in the cardiac lineage, we first evaluated bulk RNA-seq data from mESCs and CPCCs (Extended Data Fig.3b). Although the number of RaMA genes in the two cohorts is comparable, the gene sets are distinctive. The mESCs and CPCCs are heterogenous (Fig.2b, c), thus making it difficult to accurately assess RaMA gene expression. Therefore, to investigate RaMA establishment in homogenous cell populations, we next evaluated scRNA-seq data of mESCs, CSCs and CPCs. Contrary to bulk-RNA-seq data, we observed modest increased establishment (1.44%, 2.8%, 2.65%, respectively) of RaMA gene expression within the total expressed genes in mESCs, CSCs and CPCs (Fig.3b). Further, except for CMs, other cardiac cells also exhibit an increase in the number of RaMA genes. This is perhaps because the mouse CMs are multinucleated and therefore, when analyzing the CMs at the single cell level, but not on the single nucleus level, the true representation of RaMA genes in CMs has not been captured. Next, we assessed whether RaMA gene establishment in CSCs and CPCs is required for establishing DeMA genes in more differentiated cells of the cardiac lineage. However, we found that these two instances are independent processes (Extended Data Fig.3c) and that RaMA genes are mostly cell type specific, thus RaMA gene establishment may not directly correlate with lineage commitment (Extended Data Fig.3d).

### RaMA gene regulation is distinctive

Expression from a single allele is common to both RaMA and DeMA genes. Therefore, we speculated that the active and inactive alleles of these two gene cohorts may share similar histone signatures. Active RaMA gene alleles are primarily marked by H3ac, H3K27ac, H3K4me3, H3K79me2/3 and H3K9ac histone signatures (Fig.3c, Extended Data Table 2). H3K27ac is known to mark active gene enhancers^23, 24^. H3K4me3 and H3K9ac and; HeK27ac and H3K79me2 bivalently occupy highly active gene promoters^25^. It has been hypothesized that once established, active alleles and silent alleles are ‘bookmarked’ in a clonal cell population^40^. H3K79me2/3 is known to establish at active gene-bodies in an *in vitro* and *in vivo* CM differentiation paradigm^41^. The levels of H3K79me2 are known to fluctuate but are not erased during mitosis^42, 43^, thus H3K79me2/3 may serve as a ‘bookmark’ of the active allele of a RaMA gene. However, in our analysis, we found H3K36me3 as the only histone signature associated with heterochromatic RaMA genes. H3K36me3 is known to recruit HDAC to prevent redundant run-off RNA Pol II transcription^44, 45^ thus, a loose-interplay between H3K36me3 and H3K79me2 in the allele body may explain a role in orchestrating alleles to gain their active or silent signatures in a clonal cell population. Further, H3K36me3 and H3K79me2 marks have been shown to be present in nucleosomes and H3K36me3 is enriched within exons, which has been suggested to inhibit pre-mRNA splicing^46–48^. We found that most of the RaMA genes express five to six alternatively spliced transcripts on average (Fig.3d), however, 64% to 74% of these genes result in a single identified protein coding transcript (Fig.3e), suggesting a possible histone model in which H3K36me3 may safeguard the unnecessary splicing events in the single protein coding transcripts of the RaMA genes (Fig.3f). Interestingly, ENCODE histone modification data for our RaMA gene set from the six cell types we studied reveals greater similarity to maternal DeMA genes, but is distinctive from pat-mono histone signatures (Extended Data Table 1 & 2). However, at this point, we are unable to provide an explanation for this observation, as ours or others’ published data do not lead to any avenue to explore the above observation.

To investigate whether the histone signature model we proposed for RaMA genes by in-silico analysis could be validated by experimental evidence, we evaluated our CUT&Tag data for a RaMA gene histone signature. Because the number of RaMA genes we observed in our bulk-RNA-seq data from mESCs was low (13 genes), we used the RaMA genes from scRNA-seq data in mESCs. We found 85.6% of the RaMA genes agreed with the enriched promoter and gene body histone signatures in our CUT&Tag data (Extended Data Fig. 3e-g), indicating the validity of the histone signature model we proposed for RaMA genes.

### DeMA genes’ critical impact on the heart

Some DeMA genes are common to CSCs and CPCs, but some are unique to each cell type (Fig.4a). Because of the lineage relationship of these cells, we sought to assess the significance of common and unique monoallelic genes in heart development. Surprisingly, we found that unique pat-mono alleles in CPCs are exclusively regulated by 10 TFs (Extended Data Table 3); among this group five are Zinc finger proteins (Zfps) - (Fig.4b, Extended Data Fig.4a). Further, those 10 TFs are known to be involved in the processes of muscle and heart cell homeostasis (Fig.4c, Extended Data Table 4). A significant number of uniquely expressed mat-mono and pat-mono gene cohorts in CSCs and CPCs are enriched for processes related to heart development (Extended Data Table 4); noticeably, mat-mono genes contribute to embryonic heart development and young and adult heart cell physiology (e.g., MGS206: P=2.04E-10, MGS210: P=5.92E-10 - Fig.4d, Extended Data Fig.4b), while pat-mono genes are involved in young and neonates’ heart musculature (e.g., MGS844: P=0.01, MGS766: P=0.037 - Fig.4e). The genes common in CSCs and CPCs in pat-mono and mat-mono cohorts exhibit diverse and broad functional profiles (Extended Data Fig.4c, d); mat-mono genes are significantly involved in cardiomyocyte physiology (e.g., MGS320: P=3.78E-18, MGS328: P=3.74E-17), whereas pat-mono genes (e.g., Kcnd3 - cardiac repolarization) participate in heart and skeletal muscle physiology (e.g., MGS1392: P=0.014, MGS414: P=0.015) (Extended Data Fig.4e, f, Extended Data Table 4). The number of mat-mono genes that were common in four cell types from CBs was notably higher than the number of pat-mono genes (Fig.4f, g) and some (e.g., *Coa5*, *Ndufs1*, *Ndufa10*) were critical for known cardiac related human disease phenotypes (e.g., Hypertrophic cardiomyopathy- HP:0001639 – Padj= 9.942x10^-4^) (Fig.4h), whereas pat-mono genes were not (Extended Data Table 4). CMs are the most specialized cell type in the heart and perform rhythmic beating, the primary function of the heart. We assessed the importance of the unique DeMA genes in CMs in disease causation. Unique mat-mono and pat-mono genes exhibited a wide functional profile (Extended Data Fig.4g, h, Extended Data Table 3). However, the significantly higher TF functional profile for unique mat-mono genes compared to pat-mono genes suggests that the mat-mono genes hold greater risk factors for heart disease in males (Fig.4i, Extended Data Fig.4i-k, Extended Data Table 3). Conversely, unique pat-mono genes in CMs do not provide insight into known human disease related phenotype or physiological relevance.

**Fig. 4.**
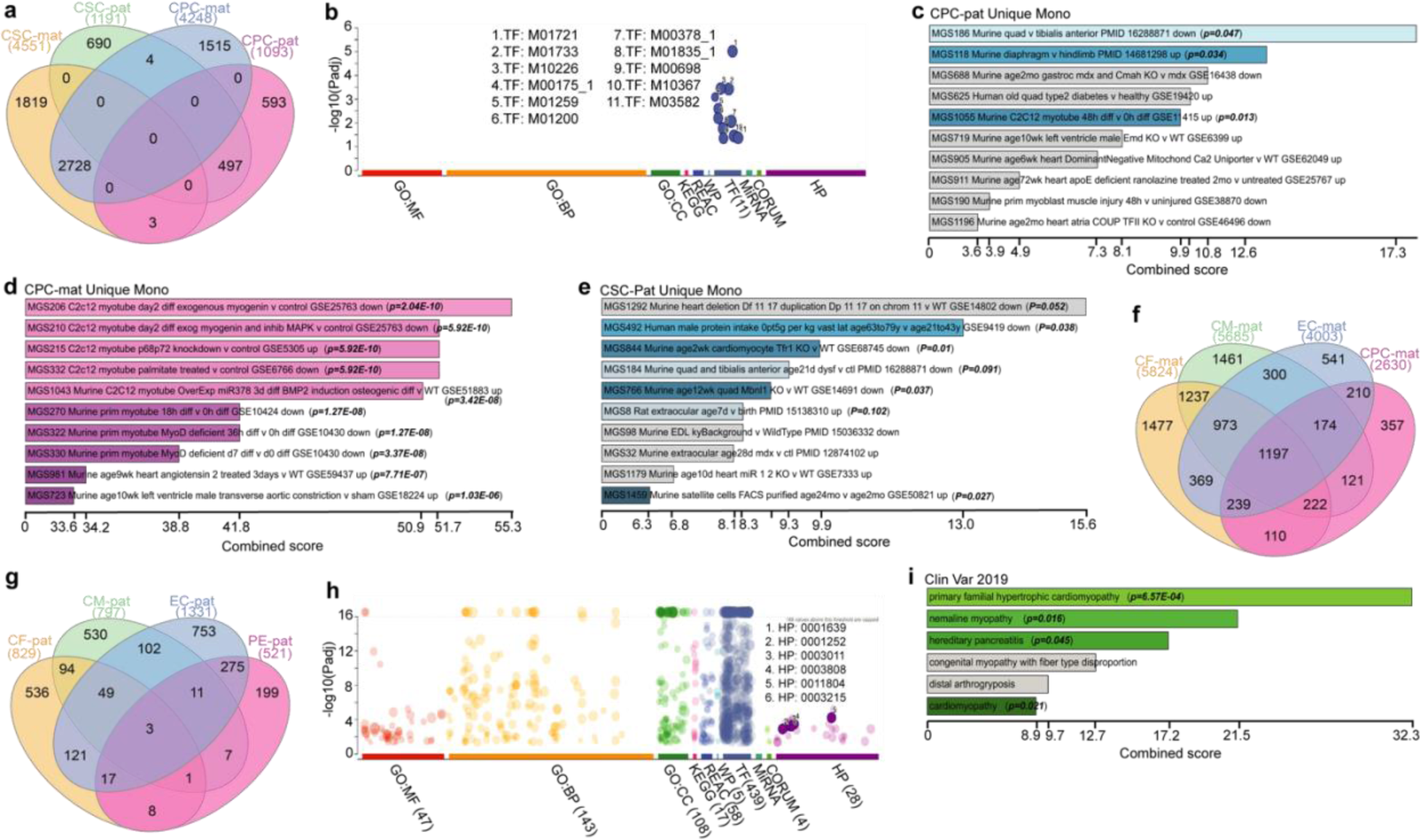
Critical cardiac genes are DeMA. **a**, Intersects of mat-mono and pat-mono genes in CPC and CSC cells. **b**, Functional profiles of CPC pat-mono genes. 588 out of 592 CPC pat-mono genes are only enriched in TF functional profile enrichment category. Data precented as combined scores (x axis) and p-value for each term. **c**-**e**, Unique CSC and CPC maternal and paternal monoallelic genes enrichment in known cardiac muscle development gene sets. Each unique monoallelic gene set was compiled with SysMyo muscle gene data and data is presented as combined scores. The length of the bars represents the combined score for each enrichment term. **f,g**, Intersects of mat- mono (**f**) and pat-mono (**g**) genes in cells from CBs. Greater number of mat-mono genes are shared between all the cell types. **h**, Functional profiles of mat-mono genes common in four cell types in CBs. Highest enrichment scored for TFs. Only the cardiac specific human phenotype IDs are shown. **i**, Disease causality with CMs’ unique mat-mono genes in the heart. Disease corelations were curated from Clin Var 2019 data. Data precented as combined scores.

To gain insight into DeMA and RaMA genes’ functional relevance in CMs, we then assessed the functional profiles of these two gene classes. Gene ontology profiles showed that the DeMA genes were enriched in processes contributing to CM cell composition (85 - GO: terms), whereas RaMA genes exhibit no association with CM structure (Extended Data Fig.4l, m). Further, only the DeMA genes exhibit enrichment in molecular function (62 - GO:MF terms), biological processes (259 - GO: BP terms) and transcription factor (639 - GO: TF terms) GO terms in regard to CMs. Moreover, from the total DeMA genes in CMs, 718 genes homologous to human were known to cause abnormalities in human heart musculature (GO: HP:0003011) (Extended Data table 5). Interestingly, we also found that, out of 718 genes, 69.1% of genes were included in autosomal recessive inheritance genes in human (HP:0000007), confirming the significance of DeMA genes in heart musculature. However, there was no strong evidence of the involvement of RaMA genes in the establishment of cardiac musculature. RaMA genes were generally related to basic cellular housekeeping processes, for example *Smug1*: base excision repair, and *Lig3*: DNA ligase (Extended Data Fig. 4n, o, (Extended Data table 6). We further investigated whether there was any known clinical information associated with DeMA genes. By compiling our DeMA gene set (6284 genes) in ‘Jensen DISEASE’, an algorithm for clinical literature data mining, we found that 28 genes are included out of 62 clinically known congenital heart disease related genes (Extended Data Fig. 4p, q, Extended Data table 7 and 8).

## Discussion

Monoallelic gene expression by imprinting is a well studied phenomenon and significantly contributed to our understanding of gene expression in development and disease^41, 49, 50^. However, it has also been shown that, for some genes, monoallelic gene expression could be explained by non-imprinting mechanisms^51^, for example establishment of recessive and dominant alleles^52^ of which the molecular mechanism is poorly understood. Also, the imprinted status of some genes may not be stable across multiple tissues, which could arise from the tissue context^53, 54^ - https://www.geneimprint.com/site/genes-by-species. In this study, using our data from mESCs and publicly available data from mESCs and mouse blastocysts, we identified a role for genetic background to influence monoallelic gene expression, which may influence the establishment of dominant and recessive alleles because of the higher polymorphisms between the two genomes. mESCs and cells in blastocysts are naïve cells and contain unique genetic and epigenetic marks, for example, gene imprinting is less abundant in those two cell types compared to lineage committed cells. Consistent with this, Rivera-Muilia et al. demonstrated that the higher number of events of asynchronous DNA replication between alleles is highly correlated with the unorganized nuclear architecture of naïve mESCs, but was lost in differentiated cells with structured nuclear architecture^55^. Therefore, the genetic background of the parental allele’s influence on deterministic genes’ allele selection (active vs inactive) is perhaps due to mESCs unique genetic and epigenetic status and may not follow the “rules of stable monoallelic gene establishment” as in lineage committed cells. To support this idea, we showed that the parental allele identity of the known imprinted genes (*Snrpn*, *Cobl*, *Dlk1*, *Meg3* and *Ubea3a*) match with our CMs DeMA data. This observation suggests that, imprinted genes may not be strongly influenced by the genetic background of the parental alleles, but the non-imprinted monoalleleic genes that are established by other mechanisms, for an example dominant and recessive gene establishment, may change the origin-of-the active allele depending on the genetic background. Therefore, we intentionally avoided categorizing the DeMA genes identified in this study as imprinted genes, but rather broadly acknowledge them as deterministic monoallelic genes. Although we were not able to address this finding further as it is beyond the scope of our study, it will be worth pursuing in future studies using the lineage committed cardiac cells from reciprocal crosses at single cell resolution.

Our data demonstrates the potential distinctive gene regulatory mechanisms in autosomal monoallelic maternal and paternal genomes in cardiac lineage cells, emphasizing monoallelic gene expression and its potential effects in cardiac development and disease. An interesting and unexpected outcome of our study is the disease susceptibility based on the exclusive expression of the maternal or paternal allele. When a critical gene is expressed solely from the single maternal allele, males are at a disadvantage in acquiring variable penetrance of disease (e.g., Fig.4i – e.g., HP:0003215 – Discarboxylic aciduria). It is known that males are more susceptible to heart-related disease^56–58^. We observed that the maternally expressed set of genes in our cardiomyocyte’s DeMA cohort, which are homologous to human genes (GO term: HP0000007 - *AUH, DNAJC19, OPA3*), are linked to the male specific 3-Methylglutaconic aciduria type 2 (BTHS syndrome) disease - **Error! Hyperlink reference not valid.**. Therefore, our study provides insightful genetic evidence of maternally expressed monoallelic genes potentially being causative of male heart-related diseases. Further, exclusively expressed paternal deterministic monoallelic genes (in CPCs) primarily showed a contribution to neonatal and adult heart physiology and this gene cohort was exclusively regulated by TFs (e.g., *Esr1*, *SP1*, *Klf4*), and are known to be involved in preventing heart attack and cardiac hypertrophy^25, 33^. Further, the DeMA genes are scattered throughout the genome, however, some showed clustering, and thus may be susceptible to disease caused by locus heterogeneity^59^. In summary, in this study we emphasize the importance of allele-specific gene expression in development, homeostasis and disease and provide a comprehensive identification of deterministic monoallelic genes in unique cardiac cell types.

## Methods

### Generation of C57BL/6J-CAST/EiJ F1 Hybrid mESCs

Four week old C57BL/6J female and four week old CAST/EiJ male mice were purchased from Jackson Laboratories. C57BL/6J female mice were hormone treated and paired (five pairs) with the male CAST/EiJ mice (two females per male) and the plug rate was 30% from the matings. At 3.5 dpc, blastocysts were collected from three plugged CAST/EiJ females. In total 37 blastocysts were placed onto feeder (Applied Stem Cell, Cat. No. ASF-1014) monolayer cultures (2i medium - 15% FBS (Millipore, Cat. No. ES-009-B), 1X Glutamine (Millipore, Cat. No. TMS-002-C), 1X Non-Essential AA (Millipore, Cat. No. TMS-001-C), 0.15 mM 2-Mercaptoethanol (Millipore, Cat. No. ES-007-E), 100 U/ml Penicillin-Streptomycin (Gibco, Cat. No.15140-122), 100 U/ml Lif (Millipore, Cat. No. ESG 1107), 1 μM PD0325901 (Sigma Aldrich, Cat. No. 444968), 3 μM CHIR99021 (Sigma Aldrich, Cat. No. 361571) in Knockout DMEM (Gibco, Cat. No. 10829-018) in 96-well plates. Cultures were maintained for 14 days to establish hybrid F1 mouse embryonic stem cells (mESCs). From 37 cultures, eight mESC-like lines were selected and transferred into feeder free 2i+LIF cultures. Cells were passaged twice and *Zfp-1* expression (fwd:5’-CTCA TGCTGGGACTTTGTGT-3’, rev:5’-TGTGTTCTGCTTTCTTGGTG-3’) in each clone was assessed prior to selecting positive (male) clones. Further, from the selected three male mESC clones (sexes were further verified in RNA-seq analysis), three single cell derived clones were established for further experiments. Preliminary, mESCs were characterized by qRT-PCR for *Pou5f1* (fwd:5’-TGTGGACCTCAGGTTGGACT, rev:5’-CTTCTGCAGGGCTTTCATGT), *Nanog*(fwd:5’-CAGATAGGCTGATTTGGTTGGTGT, rev:5’-CATCTTCTGCTTCCTGGCAA) and *Gata6* (negative control, fwd:5’- CAGCCCACGTTACGATGAACG, rev:5’-AAAATGCAGACATAAC ATTCC). By performing RNA-Seq analysis, the mESCs clones were re-confirmed as mESCs. Moreover, by performing the qRT-PCR followed by Sanger Sequencing using primers which cover known single nucleotide polymorphism (SNP) in CAST/EiJ strains, the mESCs were further confirmed for hybrid genome. Allele specific analysis of the RNA-Seq data reported, on average, 5-6 SNPs per transcript.

### F1 mESC *in vitro* culture

Male mESCs were used in all the experiments. Plastic-bottom tissue culture dishes were coated with 300 μl 0.1% gelatin (Millipore, Cat.no. ES-006-B) and incubated for at least 5 minutes at 37° C prior to cell seeding. Cells were plated at 1 x 10^5^ per well in a 6-well plate (FALCON, Cat.no. 353046) clonal density in 2i culture medium (15% FBS (Millipore, Cat. No. ES-009-B), 1X Glutamine (Millipore, Cat. No. TMS-002-C), 1X Non-Essential AA (Millipore, Cat. No. TMS- 001-C), 0.15 mM 2-Mercaptoethanol (Millipore, Cat. No. ES-007-E), 100 U/ml Penicillin-Streptomycin (Gibco, Cat. No.15140-122), 100 U/ml Lif (Millipore, Cat. No. ESG 1107), 1 μM PD0325901 (Sigma Aldrich, Cat. No. 444968), 3 μM CHIR99021 (Sigma Aldrich, Cat. No. 361571) in Knockout DMEM (Gibco, Cat. No. 10829-018)). Cells were passaged when the cultures were 70-80% confluent. The cells were washed twice with 1X PBS (Thermo Fisher, Cat. No. 14190-250) and incubated for 3 minutes at a 37° C humidified incubator with 200 μl of TrypLE Express Enzyme (Thermo Fisher. Cat. No. 12605028). The enzyme was neutralized by adding 3 ml of 10% FBS (Millipore, Cat. No. ES-009-B) in Knockout DMEM (Gibco, Cat. No. 10829-018) and centrifuged at 1010 rpm for 5 minutes at room temperature to remove the supernatant. The cell pellets were replated in 2.5 ml 2i per well (6-well plates). To cryo-freeze F1 mESCs, 70% - 80% confluent cells from one well of a 6-well plate were dissociated to obtain single cell suspension and then resuspended in freezing medium (10% DMSO (Sigma Aldrich, Cat. No. D2650), 20% FBS (Millipore, Cat. No. ES-009-B) in Knockout DMEM (Gibco, Cat. No. 10829-018)) and frozen and stored at -80° C.

### Cardiac lineage cell differentiation

Male mESC cells were resuspended in differentiation medium containing 15% FBS (Millipore, Cat. No.ES-009-B), 1X Glutamax (Thermo Fisher, Cat. No. 35050061), 1X Nonessential AA (Millipore, Cat. No. TMS-001-C), 0.15 mM 2-Mercaptoethanol (Millipore, Cat. No. ES-007-E), 100 U/ml Penicillin-Streptomycin (Gibco, Cat. No. 15140-122) in Knockout DMEM (Gibco, Cat. No. 10829-018) at the 5x10^4^ cells/ml density. 20 μl drops of cell suspension (∼1000 cells/ drop) were placed on 10 cm sterile petri dish lids and turned gently upside down onto a 10 ml 1x PBS (Thermo Fisher, Cat. No. 14190-250) containing cell culture dish so that hanging drops are formed. Cells in the hanging drops were then cultured for 48 hours at 37° C, 5% CO2 cell culture incubator to form cell-aggregates. At 48 hours post-seeding, cell-aggregates were gently rinsed into a 15 ml conical bottomed FALCON tube with 10 ml of differentiation medium and the cell-aggregates were allowed to settle down at the bottom of the tube. 1 ml of direct cardiac lineage differentiation medium (differentiation medium supplemented with 0.1 mg/ml L-Ascorbic acid (Sigma Aldrich, Cat. No. A7506) – DCLDM) was gently added and then the cell-aggregates were transferred to a tissue culture petri dishe containing 14 ml of DCLDM to form embryonic bodies (EBs) (always used tipped-off P1000 filter tips for cell-aggregates transfer). Plates with floating EBs were gently shaken twice a day to avoid EBs attaching to the plate bottoms. At 72 hours post cell-aggregate plating, EBs started expressing cardiac stem cell and mesoderm lineage markers (*Nkx2.5* (fwd:5’-CCCCCAAGTGCTCTCCTG, rew:5’-CATCCGTCTCGGCTTTGT), *Gata4* (fwd: 5’- GCAGCAGCAGTGAAGAGATG, rew:5’-GCGATGTCTGAGTGACAGGA)).

### Establishment of cardiac precursor cell culture (CPCCs)

EBs at 72 hours post cell-aggregate plating were collected into 15 ml FALCON tubes, washed three times with 10 ml of 1x PBS each, without disturbing the EBs. To harvest, the EBs were allowed to settle at the bottom of the tubes. Cells in the EBs were dissociated by adding 200 μl of TrypLE Express Enzyme (Thermo Fisher. Cat. No. 12605028), incubated at 37° C, 5% CO2 cell culture incubator for 2 minutes, added 3 ml of 10% FBS (Millipore, Cat. No. ES-009-B) in Knockout DMEM (Gibco, Cat. No. 10829-018) and then mechanically dissociated by forced pipetting; up-down with P1000 pipette. Another 7 ml of 10% FBS (Millipore, Cat. No.ES-009-B) containing Knockout DMEM (Gibco, Cat. No. 10829-018) medium was added and then centrifuged at 1010 rpm for 5 minutes at room temperature. Cell pellets were then resuspended (we used EBs from one plate) in differentiation medium containing 10 ng/ml EGF (Peprotech, Cat. No. AF-100-15) and 10 ng/ml Fgf2 (Peprotech, Cat. No. 450-33) (cardiac stem cell enrichment medium – CSCEM) and plated on 0.1% gelatin (Millipore. Cat. No. ES-006-B) coated wells in 12-wells (we used EB cells from one plate per one well in a 12-well plate) plate. Cells were passaged every other day for three passages to enrich for the CPCCs. In every passaging step we used 40% of the cells from the dissociated harvested cells to re-plate. Cells from the fourth passage were used in downstream applications.

### Cardiac body (CBs) culture

48 hours post seeded cell-aggregates were cultured in DCLDM medium in a sterile petri dish for five to six days gently shaking the dishes twice a day. We combined cell aggregates from three 10 cm plates into one 10 cm plate with 20 ml DCLDM medium. DCLDM medium was refreshed every other day. Rhythmically beating non-attached cardiac bodies (CBs) could be observed by five to six days post cell-aggregate plating and were viable and functional for another four to five days (the maximum time point we tested). For the scRNA-seq experiment, beating CBs were picked by pipets using a tipped-off P1000 filter tip under a light microscope. CBs were washed twice with 1x PBS (Thermo Fisher, Cat. No. 14190-250) and then incubated with the combination of 1:1 TrypLE Express Enzyme (Thermo Fisher. Cat. No. 12605028) and Trypsin (VWR, Cat. No. 45000-666) for 5 minutes at 37° C, at the 5% CO2 cell culture incubator and then the CBs were mechanically dissociated by pipetting up-down (P1000 pipet) followed by another 2 minutes of incubation period. Finally, the EBs were dissociated into single cell suspensions by rigorous pipetting with a p200 pipette. The cell suspensions were further filtered through cell strainers (CellTrics-Partec, Cat. No. 04-004-2326) to remove the clumps.

### Single cell RNA-seq – sample preparation

The diameters of the cells from mESCs, CPCCs and CBs single cell suspensions were measured using Countess™ ll FL Automated Cell Counter (Invitrogen, Cat. No. A27974). The sizes are mESC – 17 μm, CPCCs – 16 μm, CBs – 13 μm. We observed cells from CBs sized (measured in microscopic images) between 20 μm to 25 μm in *in vitro* cultures, but they shrank when dissociated into single cells. cDNAs from single cells were synthesized using C1TM Single-Cell Reagent Kit for mRNA Seq (Fluidigm, Cat. No. 100-6201). In brief, we used 96-well C1TM Single-Cell mRNA Seq IFC-10-17 (Fluidigm, Cat. No. 100-5760) IFCs to load the cells. Once the cells were harvested cell suspensions were filtered through cell strainers (CellTrics-Partec, Cat. No. 04-004-2326) to avoid possible cell-aggregates before the cells were loaded into integrated fluidic circuits (IFCs). Approximately 750 cells were loaded onto an IFC each time and as a control External RNA Control Consortium (ERCC- Thermo Fisher, Cat. No. 4456740) RNA spike-ins were added to the lysis buffer to obtain 1:20,000 dilutions of the final cDNA samples. Cells in IFCs were processed using the Fluidigm C1™ Single-Cell Auto Prep System. After single cells were primed into compartments, cell numbers (single or multiple cells) and the quality (debris or healthy) of the cells were observed using a light microscope and recorded corresponding to each compartment. Typical total RNAs in a mammalian cell ranged from 10-30 pg. cDNAs from low yield polyA+ RNA was synthesized using SMART-Seq® v4 Ultra® Low Input RNA Kit for the Fluidigm® C1™ System (Clontech, Cat. No. 635026), SMARTer-seq® chemistry (**S**witching **M**echanism **a**t 5’ End of **R**NA **T**emplate and Locked Nucleic acid technology), compatible for Fluidigm C1™ Single-Cell Auto Prep System. cDNA from each cell was quantified using Quant- IT™ PicoGreen® dsDNA Assay Kit (Thermo Fisher, Cat No. P11496). We used 0.3 ng of dsDNA per cell, only from the high-quality cells in library preparation. Indexed libraries were generated using Illumina Nextera XT DNA Library preparation Index Kit (Illumina, Cat No. FC-131-1096 and CF-131-1002), which is capable of multiplexing (up to 96 available indexes). Indexed libraries generated for each sample from one experiment (from one IFC) were pooled together. We finally obtained 9-pooled libraries and compressed those libraries again to 8-pooled libraries. Libraries were cleaned up using Agencourt AMPure XP beads (Beckman Coulter, Cat. No. A63880) and library quality and quantities were assessed by Agilent 2100 bioanalyzer using high sensitivity DNA Analysis chips (Agilent, Cat. No. 5067-4626). All our libraries fell within 500 bp average size and concentration ranging from 6.5 nM to 12 nM. In total we obtained 215 libraries for ESC, 175 libraries for CPC and 232 libraries for CM and cardiac smooth muscle cells. We performed 125 nt paired end sequencing for these 8-pooled libraries (one pooled library per lane) in Illumina HiSeq2500 – V4 flow cell platform at the Cold Spring Harbor Laboratory Genome Center.

### Bulk RNA-seq - Sample preparation

Cells from three mESC and CPCCs biological replicates were used. Total RNAs from each sample were extracted using RNAeasy Mini Kit (Qiagen, Cat. No. 74104) including the DNA digest step according to the manufacturer’s guidelines. All the replicates were processed in parallel at each step. 0.5 μg of total RNA from each sample was used to synthesize cDNAs from polyA+ RNA using Clontech SMART-Seq® v4 Ultra® Low Input RNA Kit for the Fluidigm® C1™ System (Clontech, Cat. No. 635026) following the manufacturer’s guidelines. Quality and quantity of cDNA from each replicate was assessed using the Agilent 2100 bioanalyzer using high sensitivity DNA Analysis chips (Agilent, Cat. No. 5067-4626). All the libraries fell within 500 bp average size. We used 0.3 ng of dsDNA per replicate for library preparation. Indexed libraries were generated using Illumina Nextera XT DNA Library preparation Index Kit (Illumina, Cat No. FC- 131-1096 and CF-131-1002). We performed 125 nt paired end sequencing for each library independently on the same chip (one library per lane) in Illumina HiSeq2500 – V4 flow cell platform at the Cold Spring Harbor Laboratory Genome Center.

### Single cell RNA-seq analysis

Sequencing data from each single cell library from each experimental point was aligned to GRCm38_v97 (mm10) mouse reference genome using STAR 2.7 aligner embedded in Partek Flow software (trial version - https://www.partek.com/partek-flow/) following the guidelines to obtain gene count tables. For the trajectory analysis, we used ASAP v1 software suite^60^. Single gene count tables of 622 scRNA-seq libraries were processed using the DDRTree reduction method. For cluster analysis, we used separate gene count tables from three independent timepoints. Gene count tables were then used in the Seurat 3.1.0 (SeuratV3) single cell data analysis package embedded in Nucleic Acid Sequence Analysis Resource^61^, a web-based portal for cell clustering. In brief, for each experiment the gene expression was normalized by “LogNormalize”, setting the scale factor to 10000 to obtain the log-transformed data. To detect the variable genes across the single cells in each experimental point per se and to calculate the average expression and depression for each gene, we applied the recommended generic settings (Mean function-ExpMean, Dispersion function-LogVMR, X Low Cut-off value: 0.0125, X High cut-off value: 3 and Y cut-off value: 0.5) to mask the outliers. To regress out the variant caused by technical noise, batch effects and biological source of variations, which would affect the clustering, we relied on the nCount_RNA feature embedded in Seurat 3.1.0 package. After computing the linear dimensional reduction to determine the statistically significant PCs, we used supervised Elbow methods and performed cell clustering with the resolution set-up at the 0.6 in default. Then, the non-linear dimension reduction was proceeded with UMAP. Each cell cluster (CPC, CSC, CF, CM, EC and PE) was then defined by using the known cell type specific markers^62–65^. Identification numbers (IDs) of each cell in each cluster was then matched with the single cell library IDs in the original ‘.fastq’ file library IDs in order to isolate the single cell libraries into relevant cell types. Cell cycle analysis was performed in StemChecker web-portal (http://stemchecker.sysbiolab.eu). Deterministic monoallelic and RaMA genes positioning in the genome were performed using Circa software (https://omgenomics.com/circa/).

### Bulk RNA-seq analysis

Differential gene expression of mESCs and CPCCs bulk-seq data was assessed with DeSeq2, an R package for “Differential gene expression analysis based on the negative binomial distribution” embedded in the NASQAR web-portal^61^.

### Allele specific scRNA-seq and bulk-RNA-seq analysis

To assess the allele-specific single cell and bulk cell analysis, we used MEA pipeline^8^ because of MEA pipelines’ usage of indels together with the SNPs in the genome to call the parental background of the transcriptomic data. We manually implemented the pipeline with all its dependencies using the Cold Spring Harbor Laboratory computing cluster. In brief, mm10 reference genome (mgp.v5.merged.snps_all.dbSNP142.vcf.gz) and genetic variants (.vcf) including SNPs and INDELs (mgp.v5.merged.indels.dbSNP142.normed.vcf.gz) were downloaded from the Ensemble genome bowser (https://useast.ensembl.org/info/data/ftp/index.html) and in silico diploid genome for C57BL/6J and CAST/EiJ was reconstructed utilizing SHAPEIT2. We then aligned each RNA-seq read file (.fastq) to the in silico genome using STAR aligner (STAR- 2.5.4b). Aligned ‘.sam’ output files were then converted into ‘.bam’ files using SAMtools- 0.1.19^66^. This process generated two .bam files each for C57BL/6J and CAST/EiJ per single cell. We then assembled all the ‘.bam’ files from a cell cluster into two separate data sets each for C57BL/6J and CAST/EiJ strains. Those two ‘.bam’ files data sets were then used for differential gene expression analysis using EdgeR pipeline embedded in SeqMonk bio-informatics web-portal (https://www.bioinformatic.babraham.ac.uk/projects/seqmonk/). The gene count-tables were then filtered through a custom written R script to filter in the deterministic monoallelic (Extended Data Fig.1e) and RaMA (Extended Data Fig.3a) genes. For bulk-RNA-seq, each experimental-data point was processed in the same way as above. Graphs and diagrams were generated using the InteractiVenn^67^, Intervene shinyapp^68^ and GraphPad PRISM (version 7.04).

### Transcript number and functional protein coding gene analysis

The BioMart data mining tool (https://www.ensembl.org/biomart/martview) in the Ensembl genome browser was used to assess the number of transcripts per RaMA gene and the number of protein coding transcripts per RaMA gene. The protein coding transcripts were cross examined in the Uniprot database (https://www.uniprot.org/uniprot).

### Gene functional profile and enrichment analysis

<u>Gene functional analysis</u> was performed by using g:Profiler (version e102_eg49_p15_7a9b4d6)^69^.

Numeric IDs were put in as ENTREZGENE_ACC for the only annotated genes. For multiple testing correction the g:SCS threshold was used as the recommended method implementing 0.05 as the threshold cut-off margin. Together with Gene Ontology analysis (Molecular Function (MF), Cellular Component (CC), Biological Process (BP)), Biological pathways (KEGG, Reactome (REAC), WikiPathway (WP), regulatory motif in DNA (TRANSFAC (TF), miRTarBase (MIRNA), protein databases (CORUM) and the Human phenotype ontology (HP) were included in the analysis.

<u>Gene enrichment analysis</u> was performed using Enricher web-portal developed by the Ma’ayan lab at the Data Coordination and Integration Center at Icahn School of Medicine at Mount Sinai^70, 71^. Gene set enrichment was performed compiling the gene list with SysMyo Muscle Gene sets (created by the Duddy lab https://www.sys-myo.com/muscle_gene_sets/) embedded in Enricher. Histone association data was obtained by compiling gene sets in ENCODE histone modification 2015 database embedded in Enricher. Results were presented in a bar chart with -Log_10_(P-values), - Log_10_(adjP-values)andcombinedscores(logoftheP-valuefromtheFisherexacttestandmultiplying that by the z-score of the deviation from the expected rank ^70, 71^). Disease relevance for gene sets was assessed with ClinVar 2019, OMIM Disease, Rare Disease AutoRIF ARCHS4 Predictions and Rare Disease GeneRIF ARCHS4 Predictions data bases. Results were presented with combined scores.

### Nascent RNA and mature RNA analysis

We used the Click-iT™ Nascent RNA Capture Kit (Thermo Fisher, Cat. No. C10365) for nascent RNA capture. Triplicates of mESCs cultured on 6-well Plates were pulsed with 0.5 mM 5-ethynyl uridine (EU) for 0.5 hours in a 37° C, 5% CO2 incubator. Single cell suspensions were obtained by dissociating cells with TrypLE and cell counts were performed using the Countess™ ll FL Automated Cell Counter (Invitrogen, Cat. No. A27974). 1 X 10^7^ cells from each replicate were used to extract RNAs. Total RNAs were extracted using the RNeasy Mini Kit (Qiagen, Cat. No. 74104), including the DNA digesting step. 10 μg of total RNAs from each replicate was used as entry material in the protocol and EU incorporated RNA was precipitated overnight at -75° C. One microgram of biotinylated RNAs with 50 ml of Dynabeads® MyOne™ Streptavidin T1 magnetic beads was used to capture biotinylated RNAs per replicate and suspended the bead-RNA complex in 50 μl of wash buffer 2 and immediately processed for cDNA synthesis. We used SuperScript® VILO™ cDNS synthesis kit (Thermo Fisher, Cat. No. 11754-050) as recommended in the Click- iT™ Nascent RNA Capture protocol. Two microliters of the undiluted cDNA was used per qRT-PCR reaction. qRT-PCR was performed with PowerUp SYBER Green Master Mix (Life Technologies, Cat.no. A25743). One microgram of the total RNA from the EU pulsed RNA samples was used to synthesize control cDNA for each replicate. Nascent RNA detection primers were designed over the intron-exon boundaries, avoiding capturing transcripts with retaining introns or processed transcripts. Mature RNA detection primers were designed to expand the product over the neighboring exon-exon where the intron between is greater than 1.5 kb. Primers used were: Thbs4-mature (fwd:5’-AGGGGAACATCTCCGAAACT, rev:5’- AAAAGCGCACC CTGATGTAG), Thbs4-nascent (fwd:5’-GTGGACACCTGTGCTCTCTG, rev:5’- GTTGCAGC GGTACTTGAGGT), Ccdc80-mature (fwd:5’-ATGTTCCTCAGTTCCGATGG, rev:5’-TCCTC TCCAACACCCAAAAG), Ccdc80-nascent (fwd:5’-TGAAATTCATCGGTTGTCAG, rev:5’- ACTCCTCCAACTTCCTCTCC), Bicd2-mature (fwd:5’-AAGCTACGGAACGAGCTCAA, rev:5’-CCATGCGCAACAGAGAGTTA), Bicd2-nascent (fwd:5’-CTCAGGCTCCAGGAGAG ATG, rev:5’-CCATGCGCAACAGAGAGTTA), Hsbp1l1-mature (fwd:5’-GAACGCGGCTGA GAATCTAC, rev:5’-CATGAGGTCATCCACGTTTC), Hsbp1l1-nascent (fwd:5’-CAGCCGTA CTCCCTCAGTGT, rev:5’-CGACTTCCCATCTCTTCCAG), 4932435O22Rik-mature (fwd:5’- CTGAGGCTCTTTGGCACTTT, rev:5’-CCACAGCAGTCCTCCTAAGC), 4932435O22Rik- nascent (fwd:5’-AGCCAGCCAGCTGAACTATC, rev:5’-ATTAC CGAAGTGTCCGATGC), Mob1b-mature (fwd:5’-TCCATTCCCGAAGAATTTCA, rev:5’-GTGC CAACTCTCGTCTGT CA), Mob1b-nascent (fwd:5’-GCATCAGGGGAGCTTAAGTG, rev:5’-GTG CCAACTCTCGT CTGTCA), Tbx3-mature (fwd:5’-CCTTCCACCTCCAACAACAC, rev:5’-GCAT GCTGTTCA AATTGAGG), Tbx3-nascent (fwd:5’-CCTCCACTCCTCCAAAACAG, rev:5’-GCAG CCATG TATGTGTAGGG), Kif17-mature (fwd:5’- GCAACTACTTCCGCTCCAAG, rev:5’-CTCA CC ACCGAAGCTGTTTT), Kif17-nascent (fwd:5’-CCAGGAGTCATGGGAGTGAC, rev:5’- GCT TGGAGCGGAAGTAGTTG).

### CUT&Tag assay

To assess the allele-specific chromatin accessibilty of maternal and paternal genes in F1 mESCs, we used CUT&Tag-IT™ assay kit (Cat. No. 53160) from Active Motif^®^. mESCs were cultured in 2i medium and at 70% confluency, cells were harvested using TrypLE Express Enzyme (Thermo Fisher. Cat. No. 12605028). Cells from three wells of a 6-well plate (FALCON, Cat.no. 353046) were combined and filtered through cell strainers (CellTrics-Partec, Cat. No. 04-004-2326) to remove cell clumps. For each histone mark, 400,000 cells were used. We followed the protocol according to the assay kit manual, excluding the magnetic bead capture steps. Instead, we used centrifugation (five minutes at 600 g at room temperature per centrifuging step) to pellet the cells. Antibodies used: H3K36me3 (Active Motif - Cat. No.61102), H3K4me1 (Active Motif - Cat. No.39298), H2K27ac (Active Motif - Cat. No.39134), H3K4me3 (Active Motif - Cat. No.39160), H3K79me2 (Active Motif - Cat. No.39144), H3K79me3 (Novus Biologicals - Cat. No.NB21- 1383SS). Guinea Pig anti-rabbit antibody provided with the kit was used as the secondary antibody as well as the IgG control. We obtained ∼200,000 nuclei in the final cell suspention in every sample and the all the nuclei were used for DNA extraction. Total fragmented DNA extracted in each sample was used to construct sequencing libraries using Illumina index primers provided with the kit. Sequencing libreries were pooled (including the other two libraries from the same experiment) and sequenced in one lane using a MiSeq PE 300 V3 kit at the Cold Spring Harbor Laboratory Genome Center.

### CUT&Tag data analysis

Allele-specific and non-allele-specific data analysis was carried out using the data anlysis webtool provided by Active Motif^®^. Brifly, peak calling was performed using SEACR (Sparse Enrichment Analysis for CUT&RUN) analysis strategy which uses the globle distribution of background signal to caliberate a simple threshold for peak calling^72^. Motif calling was performed with the Homer algorithm. For allele-specific read calling we used the MEA pipeline^10^ as previously described.

### Microscopy

Rhythmically beating cardiac body brightfield movies were taken using a Zeiss Observer Z1 inverted fluorescence microscope using a 10X objective.

### Data Availability

All NGS raw data and processed files generated and used in this study have been uploaded to GEO (GSE173403). Four extended data tables and 36 source data files are submitted with the manuscript. In allele specific gene count tables each cell has two columns, each for C57BL/6J and CAST/EiJ strains. In the column headings File=cell, number=cell number and B for C57BL/6J and C for CAST/EiJ. For example: File5B means C57BL/6J alleles in 5th cell and File5C means CAST/EiJ alleles in 5th cell.

## Acknowledgements

We would like to thank members of the Spector lab for critical discussions and advice throughout the course of this study. We would also like to thank Osama El Demerdash for assistance with setting up a computational pipeline. We acknowledge the CSHL Cancer Center Shared Resources (Animal, Microscopy, Next-Gen Sequencing) for services and technical expertise (NCI 2P3OCA45508). The sequencing analysis was performed using equipment purchased through NIH grant S10OD028632-01. This research was supporrd by R35GM131833 and RO1GM042694 (D.L.S.), and grant 18-26 from the Charles H. Revson Foundation (G.I.B.).

## Author Contribution

G.I.B. and D.L.S. conceived and designed the overall study. G.I.B. performed all the experiments and analysis. G.I.B. and D.L.S. wrote and edited the manuscript.

---

**Extended Data Fig.1.**
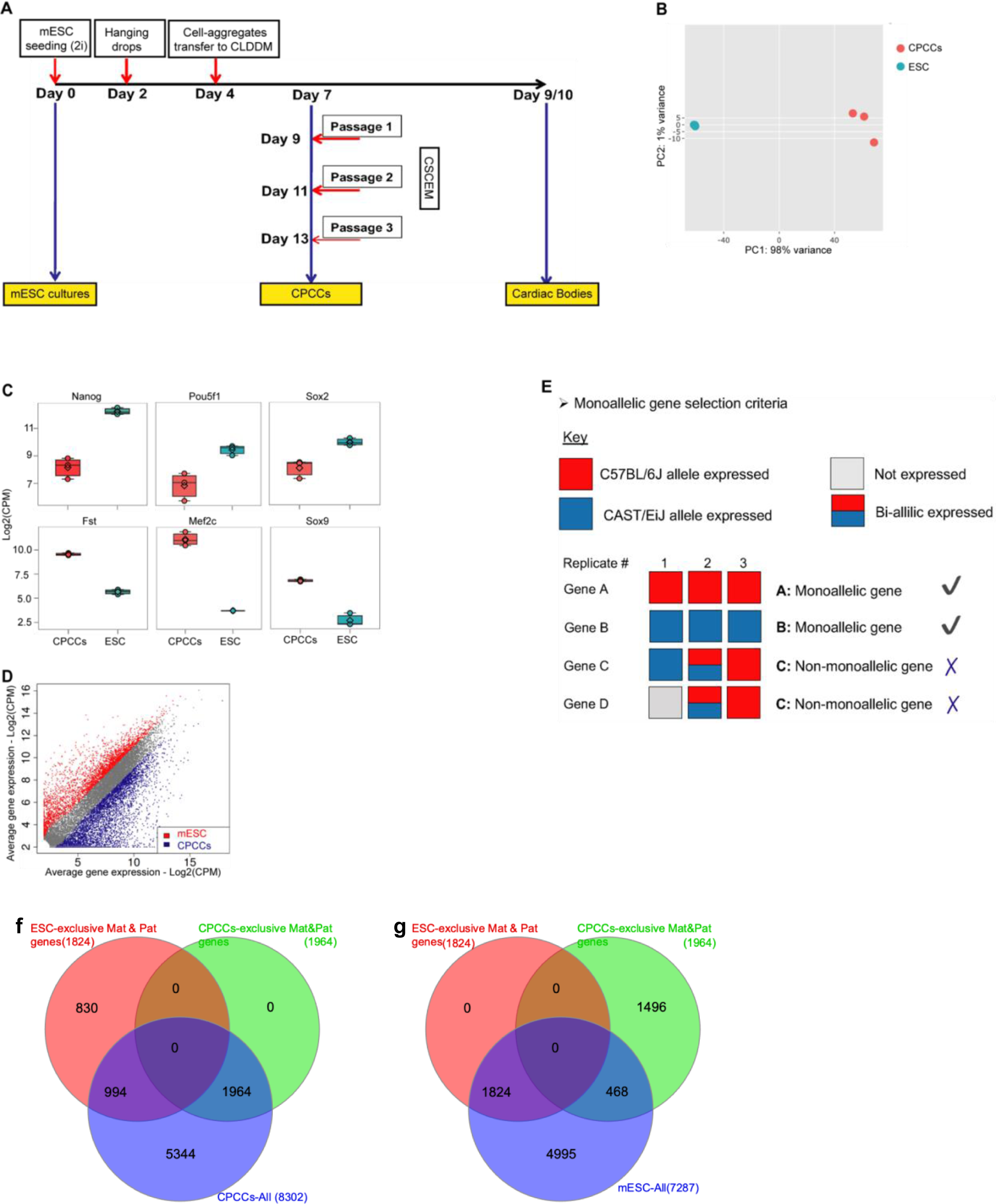
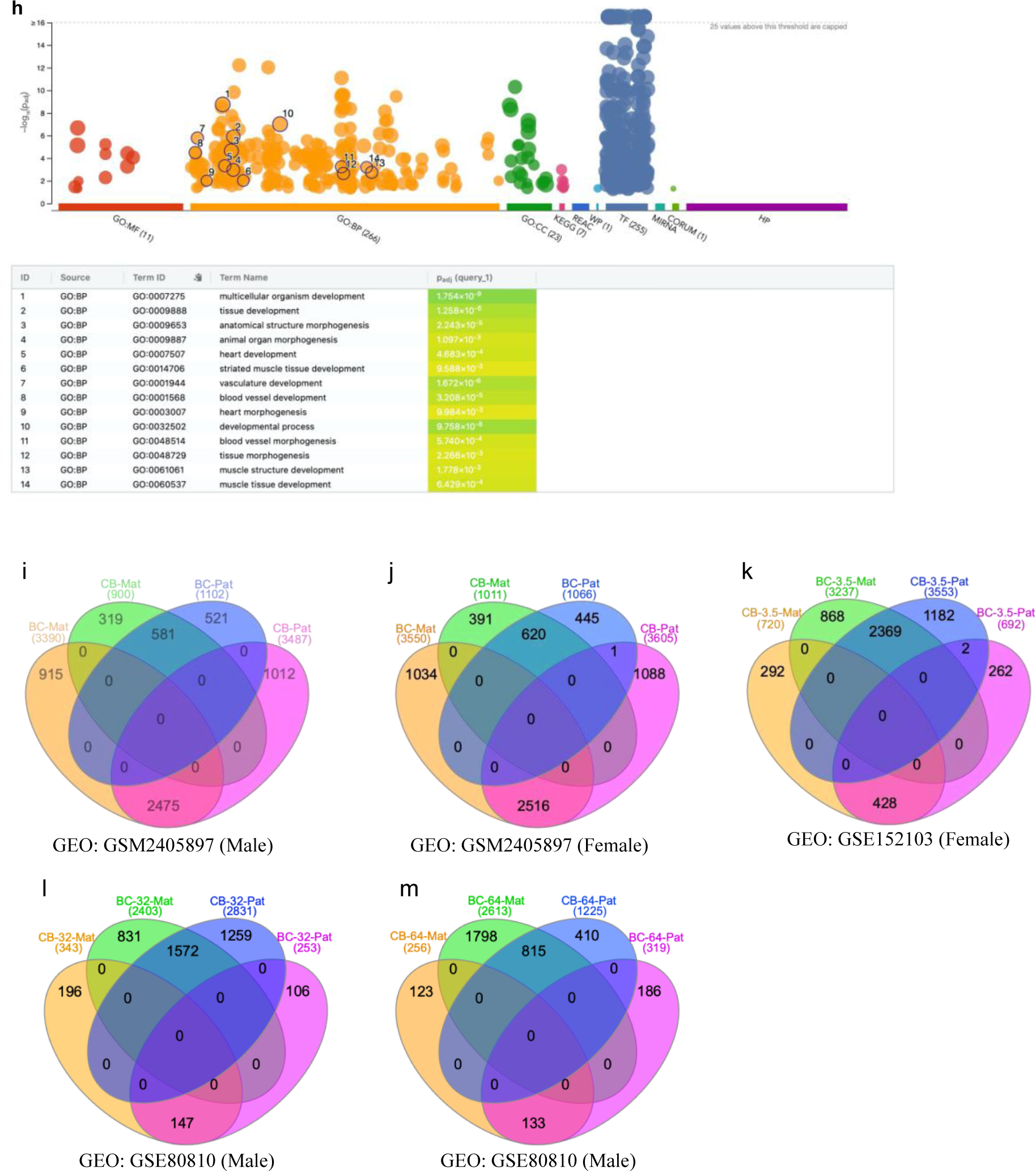
Allelic expression of autosomal monoallelic genes. **a**, Schematic of in vitro cell differentiation protocol of cardiac lineage cell types. Cardiac intermediate cultures at day 7 are highly heterogenous cell cultures. CSC and CPC enriched cultures were obtained passaging these intermediate cultures three times in enrichment medium before harvesting for RNA-seq experiments. At 9-10 days post ascorbic acid treatment, detached embryonic bodies start rhythmically beating and become cardiac bodies. Picked floating-beating cardiac bodies were used in scRNA-seq experiments. **b**, Unsupervised Principal Component Analysis (PCA) of mESCs and CPCCs bulk-RNA-seq data. **c**, Expression levels of known mESC and CSC-CPC marker genes. **d**, Differential gene expression in mESCs and CPCCs. Differentially expressed mESC and CPCCs genes are uniquely enriched in each cell type. -Log2(2) CPM was uses as the cut off. **e**, Schematic of the deterministic monoallelic gene selection criteria. Genes are categorized as: not-expressed, maternal-allele only expressed, paternal-allele only expressed and bi-allelically expressed. To be categorized as a monoallelic gene (1), the gene must be expressed in all biological triplicates, and 1. (2) the gene must be exclusively expressed from only one parental background. **f**,**g**, Relation of monoallelic gene establishment between mESCs and CPCCs. Maternal and Paternal DeMA genes in mESCs and CPCCs were assessed among the total CPCCs genes (**f**) and the total mESCs (**g**). Venn diagrams show 994 monoallelically expressed DeMA genes in mESCs changed to biallelic expression in CPCCs and 468 biallelically expressed genes in mESCs changed to monoalleleic expression. **h**, Functional analysis of newly established DeMA genes in CPCCs. GO:BP terms clearly indicate the cardiac lineage specific functional relevance of several new monoallelic genes. **i**-**m**, Venn diagrams illustrating the effect of genetic background of the alleles in DeMA gene biased maternal and paternal expression. All the analyses show the switch of the DeMA genes depending on the genetic background of the allele. Maternal is abbreviated to “Mat” and paternal is abbreviated to “Pat”. CB: cells from CAST/EiJ x C57BL/6J crosses and BC: cells from C57BL/6J x CAST/EiJ crosses. Either in CB or BC, “C” is for CAST/EiJ and “B” is for C57BL/6J and the first letter indicates the maternal background. GSM2405897: male and female RNA-seq data was generated from mESC bulk cell samples. GSE152103: male and female RNA-seq data was generated from total cells in 3.5 dpc blastocycts. GSE80810: male and female RNA-seq data was generated from single cells of 32-cell and 64-cell blastocyct.

**Extended Data Fig. 2.**
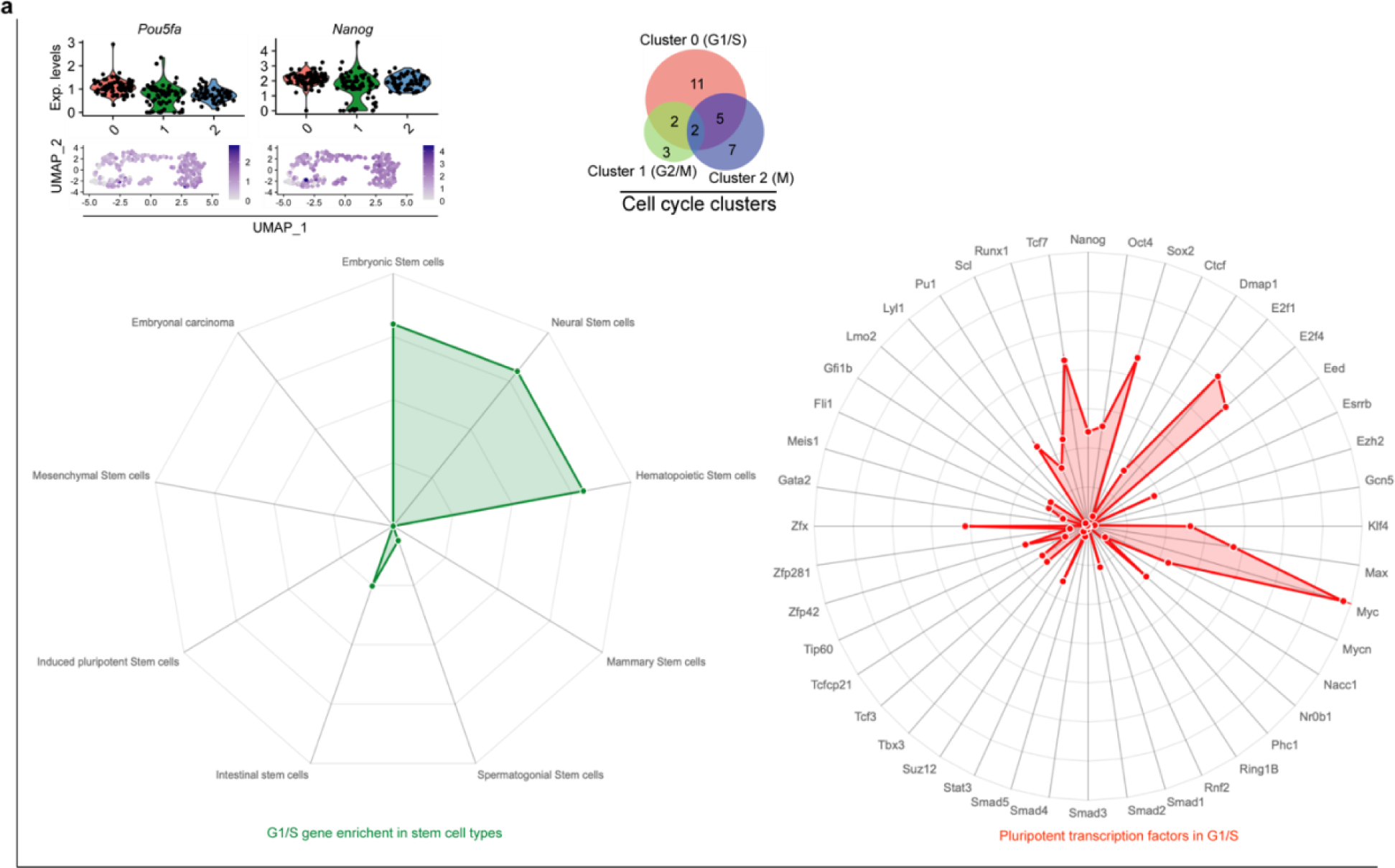
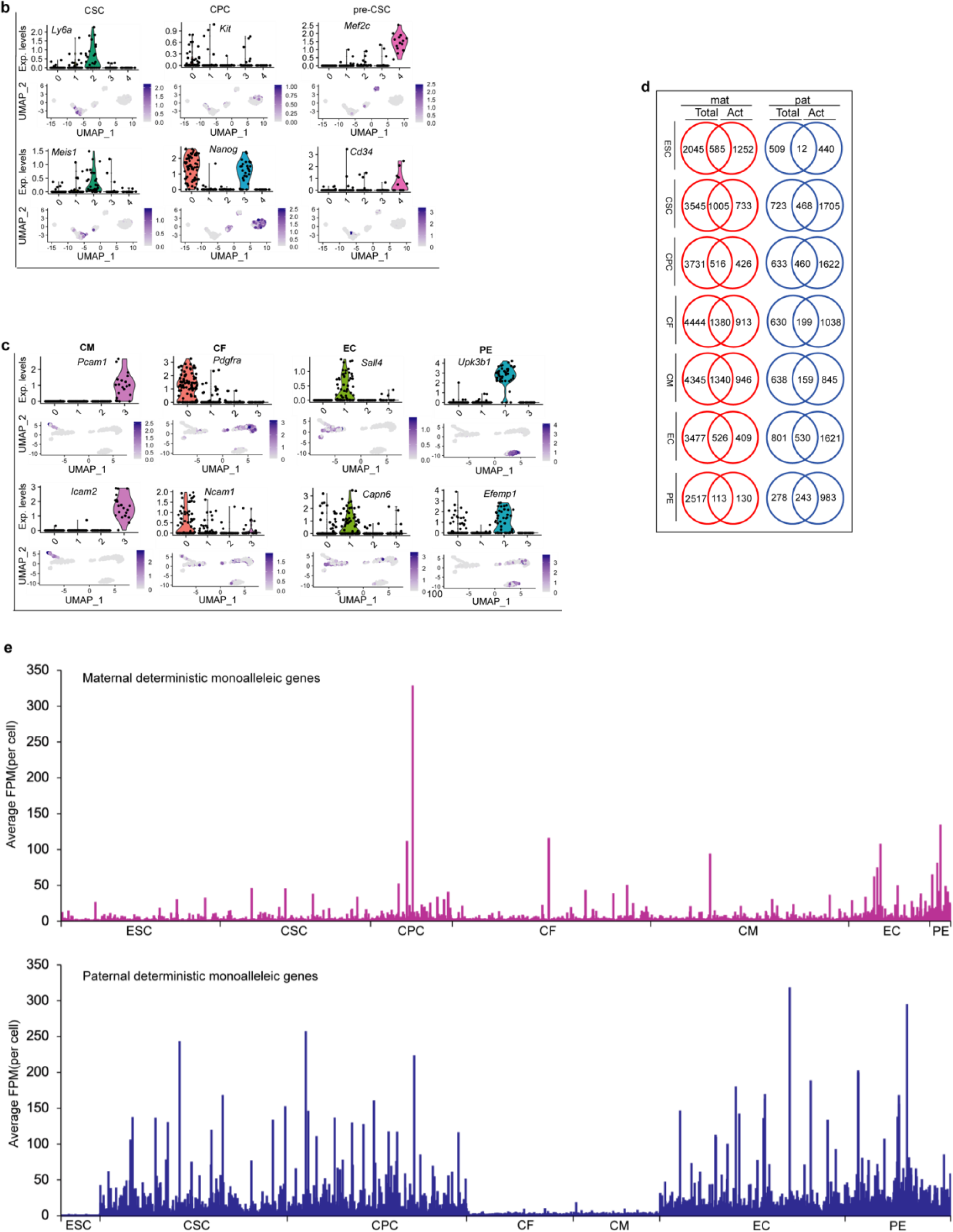
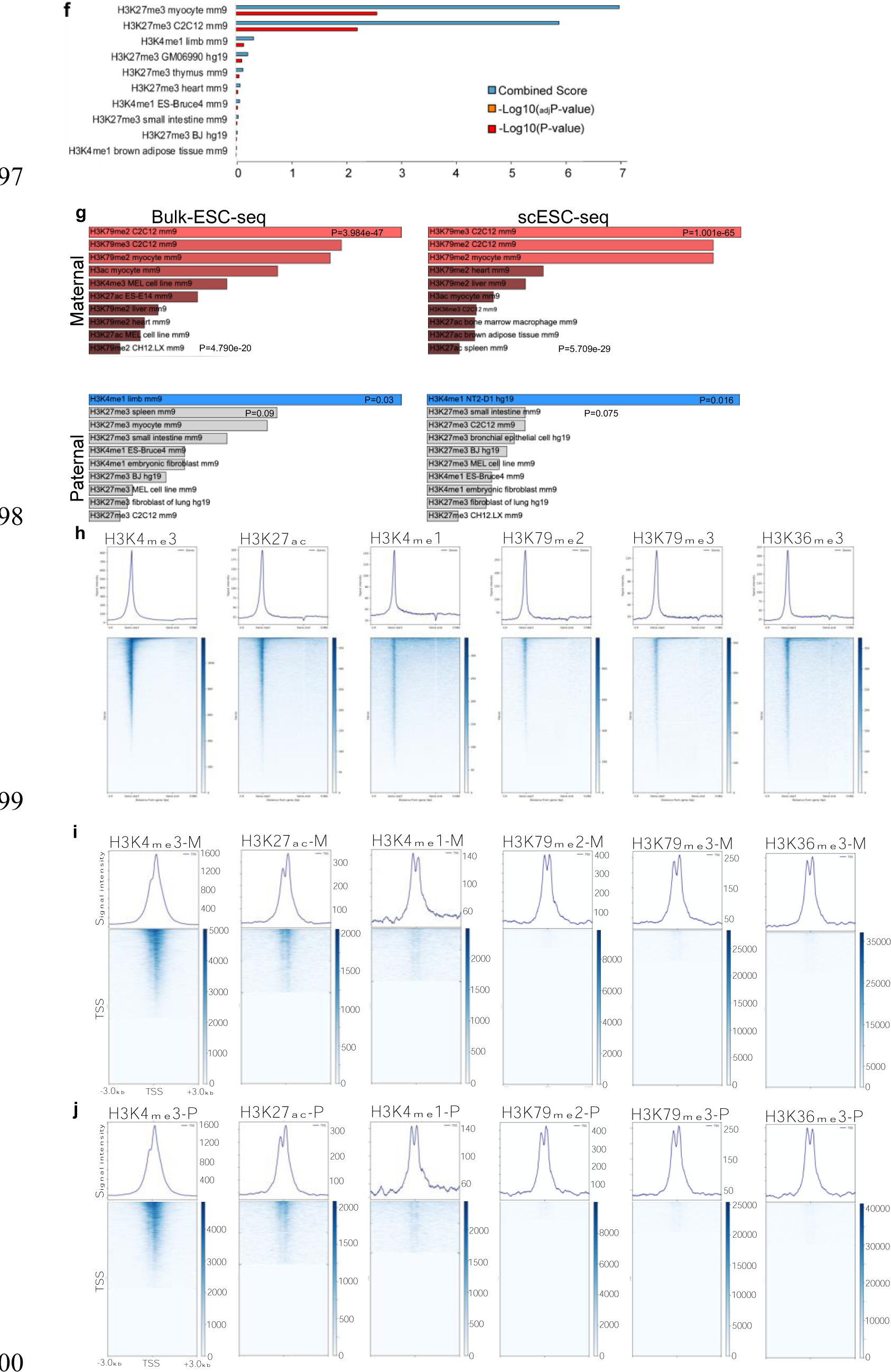
Allele expression in cardiac cell types and allele-specific differential epigenetic regulation of monoallelic genes. **a**, mESC cell clustering and cell cycle gene enrichment analysis. Gene expression levels and UMAP plots show high enrichment of *Pou5fa* and *Nanog* in three mESC clusters. Cell cycle transcriptomic analysis project the three cell clusters from three cell cycle stages: Cluster 0: G1/S, Cluster 1: G2/M, Cluster 2: M. Radar-plot showing the gene enrichment in different stem cell types (bottom-left) and key pluripotent transcription factor enrichment in G1/S cluster (bottom-right). **b**, Cell type specific gene enrichment and gene clustering (UMAP-cluster graphs) for CSC, CPC and pre-CSC cells from CPCCs cultures. **c**, Cell type specific gene enrichment and gene clustering (UMAP) for CF, CM, EC and PE in cells from CBs. **d**, Multiple Venn diagram showing the number of genes in two transcript categories, mature (Total) transcripts and nascent (Act) transcripts, for seven cell types studied. Except in mESCs and CMs, the actively expressed number of pat-mono genes is greater than the number of mat-mono actively transcribing genes. Mat-mono genes are lacking their nascent transcripts more often than pat-mono genes. **e**, Average (Fragment Per Million-FPM) maternal and paternal active gene expression for mESCs, CSCs, CPCs, CFs, CMs, ECs and PEs. Data point at the X- axis represent genes. In mESC, CFs and CMs, the transcription rates are higher in mat-mono genes and in CSCs, CPCs, ECs and PEs, the transcription rate is higher in pat-mono genes. **f**, Histone signatures of pat-mono genes. Histone marks enrichment plotted for Combined score, -Log_10_(adjP-value), - Log_10_(P-value) (see methods for statistics used). **g**, In-silico analysis of maternal and paternal DeMA genes in F1 mESCs bulk and scRNA-seq data using ENCODE histone data. For maternal genes the curated histone signatures are highly probabilistic, while for the paternal gene the probability of the histone marks curated are comparatively low p=0.03 and p=0.09). **h**, Histone signatures’ distribution throughout a hypothetical gene and intergenic regions. **i & j**, Allele specific histone association for the histone marks used in CUT&Tag assay (i-maternal and j- paternal). H3K4me3, a promoter signature indicating the presence of two peaks, a predominant one at the -TSS and a weak peak at the +TSS, most probably an indication of proximal promoters. H3K27ac and H3K4me1, enhancer signatures also indicate two peaks beside the TSS, which may also be associated with proximal enhancers at the region 5’ of the TSS. Lower number of gene’s 5’ and 3’ of TSS reagions are marked by the H3K79me2, H3K79me3 and H3K36me3 exhibiting active gene promoter association, but the majority of the enrichment is widespread across the gene bodies.

**Extended Data Fig. 3.**
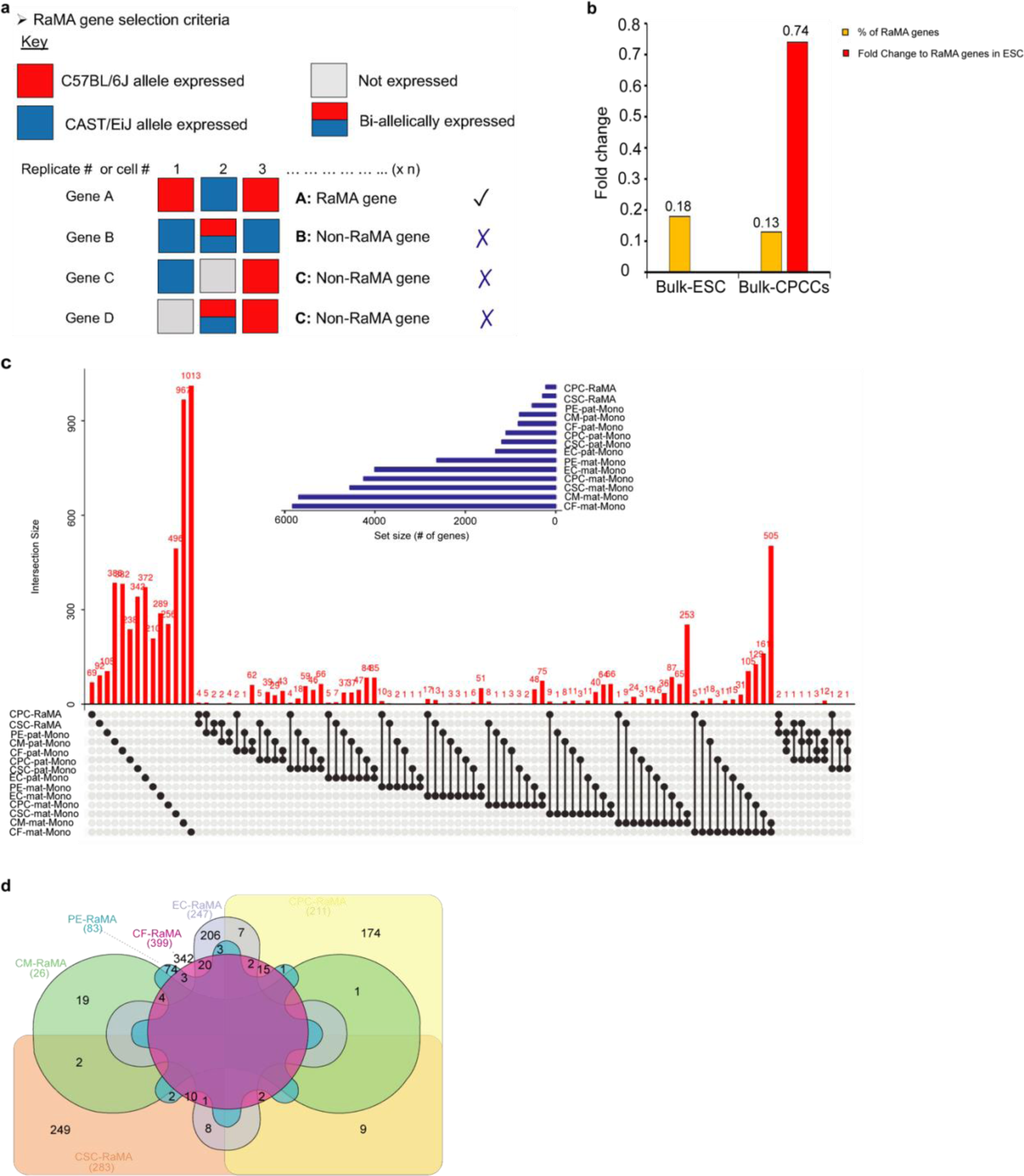
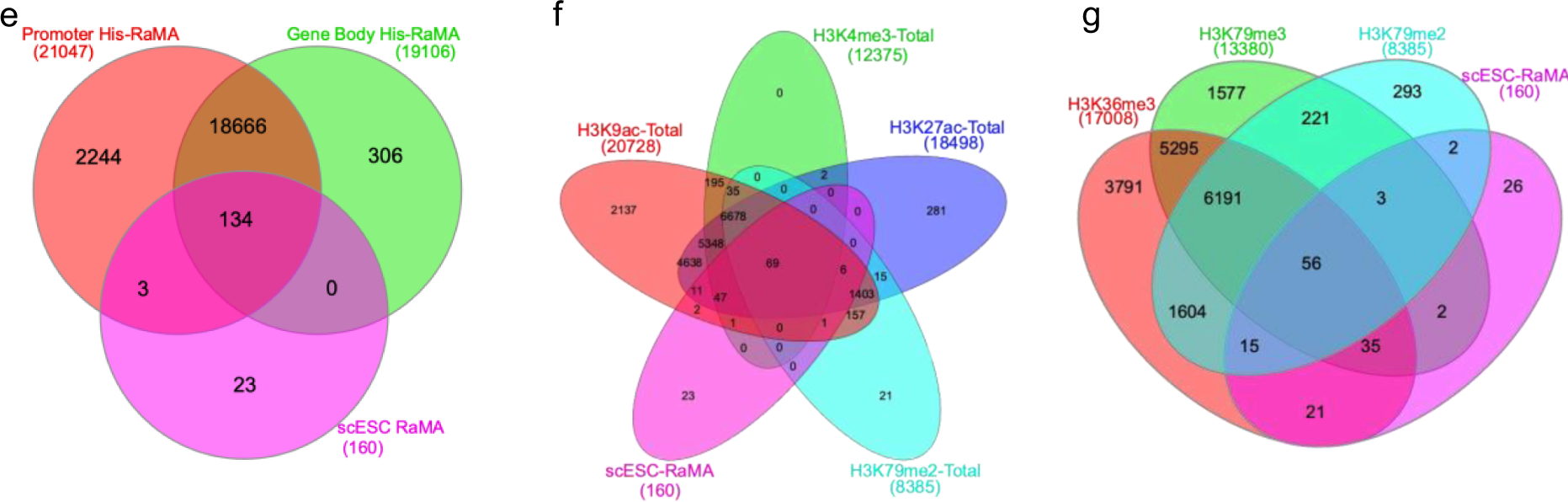
RaMA genes exhibit lineage specificity and expression is regulated by distinct histone modifications of monoallelic genes. **a,** Schematic of RaMA gene selection criteria. Genes fit into the selection criteria ‘A’ is considered as RaMA genes. **b,** Percentages and the fold changes of the expressed RaMA genes (bulk-RNA-seq). Percentages of the RaMA genes in each cells type were calculated over the total genes expressed and the fold change of the expressed RaMA genes in CPCCs was calculated over the RaMA genes from mESCs. **c,** UpSet diagram illustrationg the intersects of unique and common RaMA genes in cardiac lineage cell types. RaMA genes are preferentially unique to cell types but shared less between cell types. **d,** Intersects between CSC and CPC RaMA genes and monoallelic genes of four cell types in CBs. Nodes show the intersected gene size and the lines connecting the nodes shows the cell types that are sharing the intersected genes. **e-g**, Assesment of RaMA gene’s association with gene enriched for histone marks identified in CUT&Tag assay. 85.6% of RaMA genes (from mESCs scRNA-seq data) are associated with in-silico modeled histone signatures for promoters and gene bodies (e). Venn diagrams showing individual promoter histone signature (f) and gene body (g) association with RaMA genes.

**Extended Data Fig. 4.**
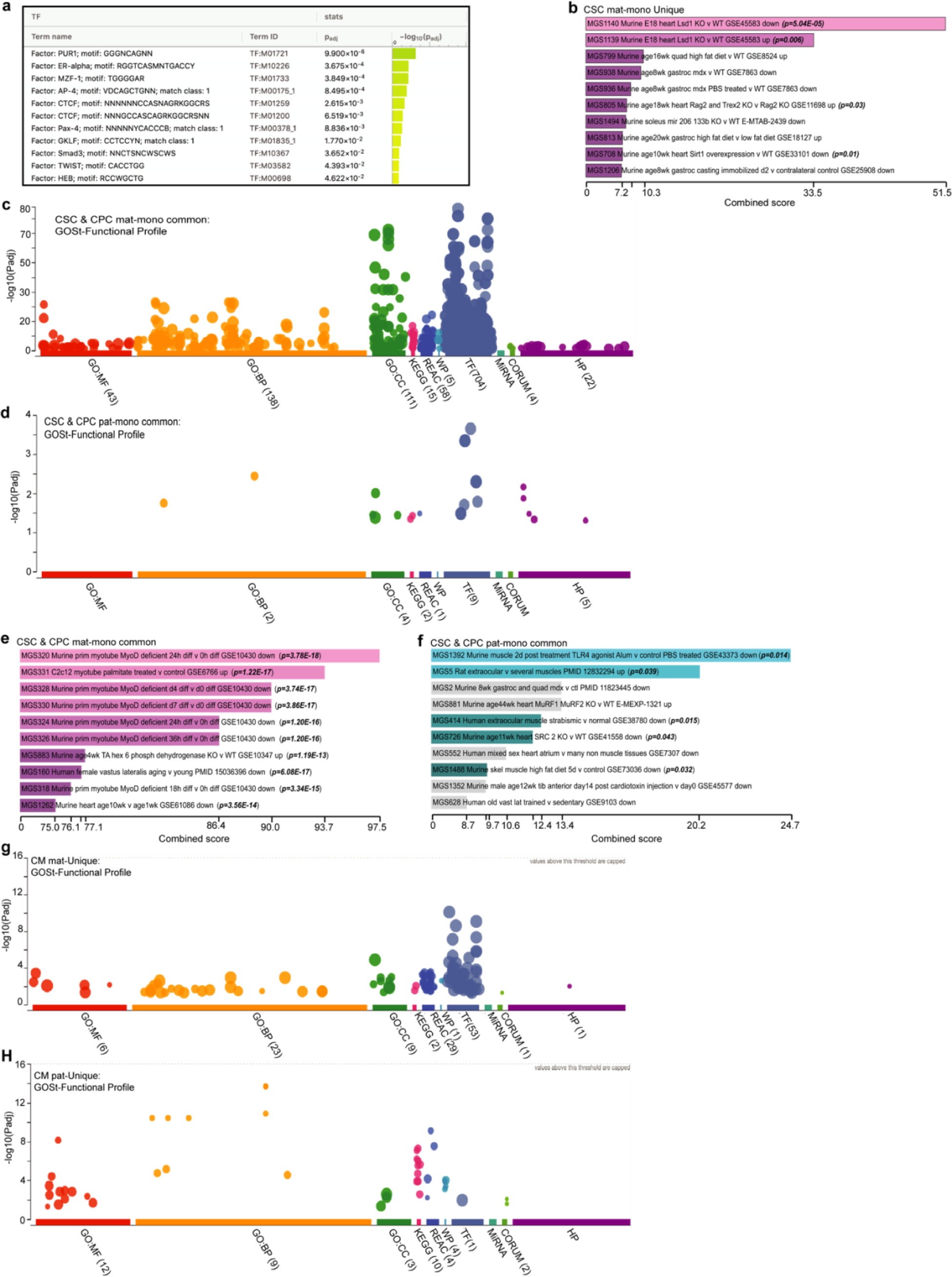
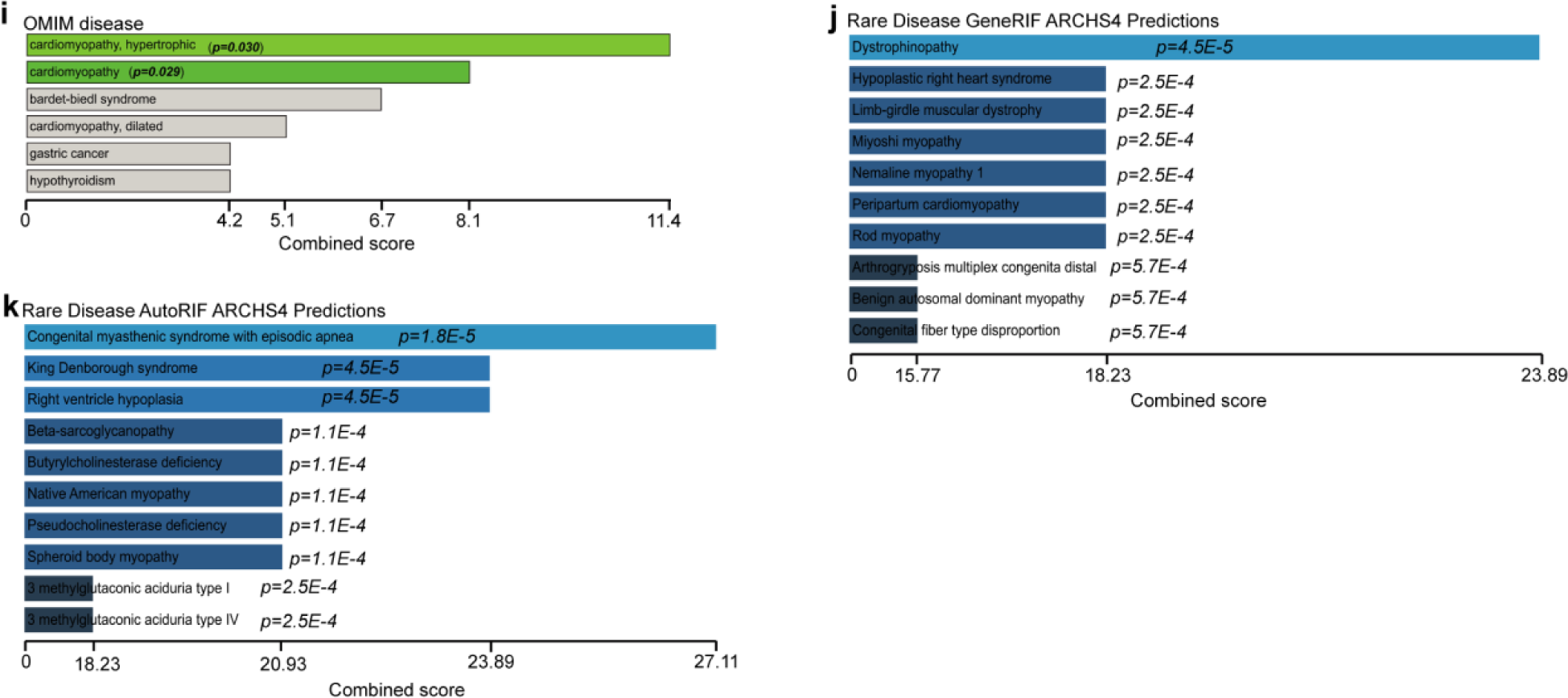
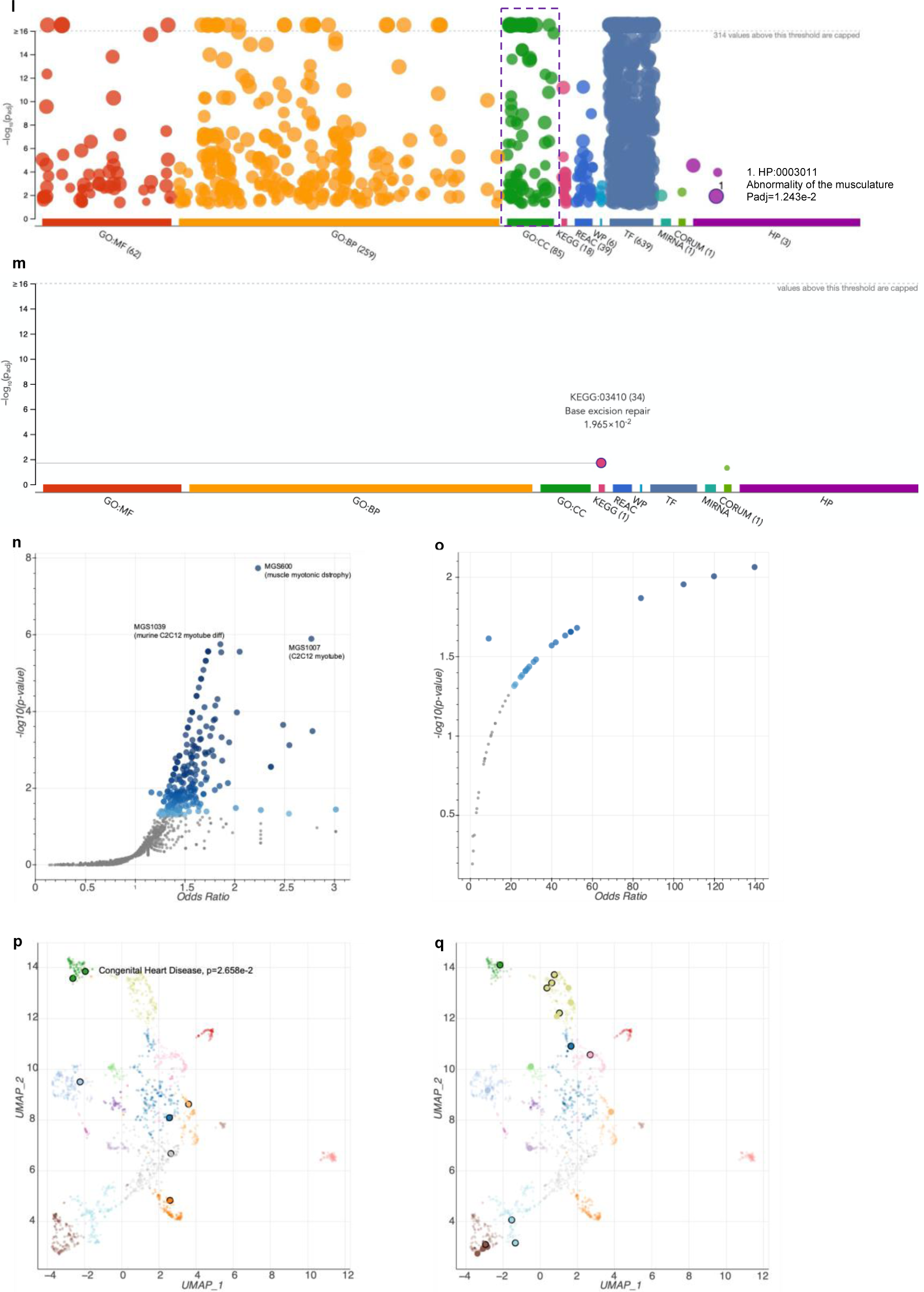
Critical cardiac genes are DeMA. **a**, Transcription factor binding motif enrichment of pat-mono genes in CPCs. GOSt profiles shows only TF binding functional profiles for CPC mat-mono genes. **b**, Unique CSC maternal monoallelic genes enrichment in known cardiac muscle development gene sets. **c**,**d**, GOSt functional profiles of CSCs’ and CPCs, common mat-mono (**c**) and common pat-mono (**d**) genes. MF, BP, CC and TF shows significantly higher GOSt profiles for mat-mono genes than pat-mono genes. **e**,**f**, Enrichment of CSCs, and CPCs, common mat-mono (**e**) and pat-mono (**f**) genes in SysMyo muscle data. Data indicates higher enrichment of common monoallelic genes in embryonic, neonatal and adult heart. **g**,**h**, Functional profiles of unique maternal and paternal monoallelic genes in CMs. Only one TF (TF:M08998) was enriched for 135 pat-mono genes. **i-k**, CMs’ mat-mono genes’ involvement in different rare disease categories. Significant involvements were curated from OMIM disease, Rare disease . Bars represent combined scores. **l**,**m**, Functional profile analysis of DeMA and RaMA genes in CMs. DeMA genes exhibi broader functional profile **(l)** than the RaMA genes **(m)** and DeMA genes are included in 85 cellular component GO terms in which there are processes included for cardiac cell composition build. Among the total DeMA gene set 718 genes (out of 1201 query size) are included in the ‘Abnormality of the musculature’ human phenotypes. **n**,**o**, Volcano plot showing enrichment of total DeMA gene and RaMA genes in CMs in ‘Sysmyo muscle data’. DeMA genes are significantly involved in cardiac cell composition related processes **(n)** while RaMA genes are involving in cellular housekeeping processes **(o)**. **p**,**q**, UMAP plots showing DeMA and RaMA gene’s known clinical relevance. The gene sets were compiled with ‘Jensen DISEASE’, a clinical literature data mining algorithm. DeMA genes are significantly (p=2.658e-2) enriched in clinical congenital heart disease.

## References

1. Deng, Q., Ramskold, D., Reinius, B. & Sandberg, R. Single-cell RNA-seq reveals dynamic, random monoallelic gene expression in mammalian cells. Science 343, 193–196, doi:10.1126/science.1245316 (2014).

2. Eckersley-Maslin, M. A. et al. Random Monoallelic Gene Expression Increases upon Embryonic Stem Cell Differentiation. Dev Cell 28, 351–365, doi:10.1016/j.devcel.2014.01.017 (2014).

3. Gendrel, A. V. et al. Developmental dynamics and disease potential of random monoallelic gene expression. Dev Cell 28, 366–380, doi:10.1016/j.devcel.2014.01.016 (2014).

4. Gimelbrant, A., Hutchinson, J. N., Thompson, B. R. & Chess, A. Widespread monoallelic expression on human autosomes. Science 318, 1136–1140, doi:10.1126/science.1148910 (2007).

5. Xu, J. et al. Corrigendum: Landscape of monoallelic DNA accessibility in mouse embryonic stem cells and neural progenitor cells. Nat Genet 49, 970, doi:10.1038/ng0617-970a (2017).

6. Naqvi, S. et al. Conservation, acquisition, and functional impact of sex-biased gene expression in mammals. Science 365, 249-+, doi:10.1126/science.aaw7317 (2019).

7. Oliva, M. et al. The impact of sex on gene expression across human tissues. Science 369, doi:10.1126/science.aba3066 (2020).

8. Richard Albert, J., et al. Development and application of an integrated allele-specific pipeline for methylomic and epigenomic analysis (MEA). Bmc Genomics 19, 463, doi:10.1186/s12864-018-4835-2 (2018).

9. Werner, R. J. et al. Sex chromosomes drive gene expression and regulatory dimorphisms in mouse embryonic stem cells. Biol Sex Differ 8, doi:ARTN 28 10.1186/s13293-017-0150-x (2017).

10. Santini, L. et al. Genomic imprinting in mouse blastocysts is predominantly associated with H3K27me3. Nat Commun 12, doi:ARTN 3804 10.1038/s41467-021-23510-4 (2021).

11. Borensztein, M. et al. Xist-dependent imprinted X inactivation and the early developmental consequences of its failure. Nat Struct Mol Biol 24, 226-+, doi:10.1038/nsmb.3365 (2017).

12. Raghavan, A. et al. Genome-wide analysis of mRNA decay in resting and activated primary human T lymphocytes. Nucleic Acids Res 30, 5529–5538, doi:10.1093/nar/gkf682 (2002).

13. Sun, W., Gao, Q., Schaefke, B., Hu, Y. & Chen, W. Pervasive allele-specific regulation on RNA decay in hybrid mice. Life Sci Alliance 1, e201800052, doi:10.26508/lsa.201800052 (2018).

14. Ono, R., et al. A retrotransposon-derived gene, PEG10, is a novel imprinted gene located on human chromosome 7q21. Genomics 73, 232–237, doi:10.1006/geno.2001.6494 (2001).

15. Carr, M. S., Yevtodiyenko, A., Schmidt, C. L. & Schmidt, J. V. Allele-specific histone modifications regulate expression of the Dlk1-Gtl2 imprinted domain. Genomics 89, 280–290, doi:10.1016/j.ygeno.2006.10.005 (2007).

16. Da Rocha, S. T., Edwards, C. A., Ito, M., Ogata, T. & Ferguson-Smith, A. C. Genomic imprinting at the mammalian Dlk1-Dio3 domain. Trends Genet 24, 306–316, doi:10.1016/j.tig.2008.03.011 (2008).

17. Tierling, S., Gasparoni, G., Youngson, N. & Paulsen, M. The Begain gene marks the centromeric boundary of the imprinted region on mouse chromosome 12. Mamm Genome 20, 699–710, doi:10.1007/s00335-009-9205-6 (2009).

18. Gendrel, A. V., Marion-Poll, L., Katoh, K. & Heard, E. Random monoallelic expression of genes on autosomes: Parallels with X-chromosome inactivation. Semin Cell Dev Biol 56, 100–110, doi:10.1016/j.semcdb.2016.04.007 (2016).

19. Levin-Klein, R. & Bergman, Y. Epigenetic regulation of monoallelic rearrangement (allelic exclusion) of antigen receptor genes. Front Immunol 5, 625, doi:10.3389/fimmu.2014.00625 (2014).

20. Inoue, A., Jiang, L., Lu, F., Suzuki, T. & Zhang, Y. Maternal H3K27me3 controls DNA methylation-independent imprinting. Nature 547, 419–424, doi:10.1038/nature23262 (2017).

21. Nag, A. et al. Chromatin signature of widespread monoallelic expression. Elife 2, e01256, doi:10.7554/eLife.01256 (2013).

22. Pathak, R. & Feil, R. Oocyte-derived histone H3 lysine 27 methylation controls gene expression in the early embryo. Nat Struct Mol Biol 24, 685–686, doi:10.1038/nsmb.3456 (2017).

23. Creyghton, M. P. et al. Histone H3K27ac separates active from poised enhancers and predicts developmental state. Proc Natl Acad Sci U S A 107, 21931–21936, doi:10.1073/pnas.1016071107 (2010).

24. Zhang, T. T., Zhang, Z. Q., Dong, Q., Xiong, J. & Zhu, B. Histone H3K27 acetylation is dispensable for enhancer activity in mouse embryonic stem cells. Genome Biol 21, doi:ARTN 45 10.1186/s13059-020-01957-w (2020).

25. Papait, R. et al. Genome-wide analysis of histone marks identifying an epigenetic signature of promoters and enhancers underlying cardiac hypertrophy. Proc Natl Acad Sci U S A 110, 20164–20169, doi:10.1073/pnas.1315155110 (2013).

26. Hyun, K., Jeon, J., Park, K. & Kim, J. Writing, erasing and reading histone lysine methylations. Exp Mol Med 49, e324, doi:10.1038/emm.2017.11 (2017).

27. Jonas, S. & Izaurralde, E. Towards a molecular understanding of microRNA-mediated gene silencing. Nat Rev Genet 16, 421–433, doi:10.1038/nrg3965 (2015).

28. Kaushik, M. et al. A bouquet of DNA structures: Emerging diversity. Biochem Biophys Rep 5, 388–395, doi:10.1016/j.bbrep.2016.01.013 (2016).

29. Bae, S. & Lesch, B. J. H3K4me1 Distribution Predicts Transcription State and Poising at Promoters. Front Cell Dev Biol 8, 289, doi:10.3389/fcell.2020.00289 (2020).

30. Chen, L. F. et al. Enhancer Histone Acetylation Modulates Transcriptional Bursting Dynamics of Neuronal Activity-Inducible Genes. Cell Rep 26, 1174–1188 e1175, doi:10.1016/j.celrep.2019.01.032 (2019).

31. Cheng, J. et al. A role for H3K4 monomethylation in gene repression and partitioning of chromatin readers. Mol Cell 53, 979–992, doi:10.1016/j.molcel.2014.02.032 (2014).

32. Reveron-Gomez, N. et al. Accurate Recycling of Parental Histones Reproduces the Histone Modification Landscape during DNA Replication. Mol Cell 72, 239–249 e235, doi:10.1016/j.molcel.2018.08.010 (2018).

33. Zhang, W. Q., Song, M. S., Qu, J. & Liu, G. H. Epigenetic Modifications in Cardiovascular Aging and Diseases. Circ Res 123, 773–786, doi:10.1161/Circresaha.118.312497 (2018).

34. Rando, O. J. Global patterns of histone modifications. Curr Opin Genet Dev 17, 94–99, doi:10.1016/j.gde.2007.02.006 (2007).

35. Pursani, V., Bhartiya, D., Tanavde, V., Bashir, M. & Sampath, P. Transcriptional activator DOT1L putatively regulates human embryonic stem cell differentiation into the cardiac lineage. Stem Cell Res Ther 9, 97, doi:10.1186/s13287-018-0810-8 (2018).

36. Kaya-Okur, H. S., et al. CUT&Tag for efficient epigenomic profiling of small samples and single cells. Nat Commun 10, 1930, doi:10.1038/s41467-019-09982-5 (2019).

37. Godfrey, L. et al. H3K79me2/3 controls enhancer-promoter interactions and activation of the pan- cancer stem cell marker PROM1/CD133 in MLL-AF4 leukemia cells. Leukemia 35, 90–106, doi:10.1038/s41375-020-0808-y (2021).

38. Godfrey, L. et al. DOT1L inhibition reveals a distinct subset of enhancers dependent on H3K79 methylation. Nat Commun 10, doi:ARTN 2803 10.1038/s41467-019-10844-3 (2019).

39. Zwemer, L. M. et al. Autosomal monoallelic expression in the mouse. Genome Biol 13, R10, doi:10.1186/gb-2012-13-2-r10 (2012).

40. Eckersley-Maslin, M. A. & Spector, D. L. Random monoallelic expression: regulating gene expression one allele at a time. Trends Genet 30, 237–244, doi:10.1016/j.tig.2014.03.003 (2014).

41. Cattaneo, P. et al. DOT1L-mediated H3K79me2 modification critically regulates gene expression during cardiomyocyte differentiation. Cell Death Differ 23, 555–564, doi:10.1038/cdd.2014.199 (2016).

42. Farooq, Z., Banday, S., Pandita, T. K. & Altaf, M. The many faces of histone H3K79 methylation. Mutat Res-Rev Mutat 768, 46–52, doi:10.1016/j.mrrev.2016.03.005 (2016).

43. Fu, H. Q. et al. Methylation of Histone H3 on Lysine 79 Associates with a Group of Replication Origins and Helps Limit DNA Replication Once per Cell Cycle. Plos Genet 9, doi:ARTN e1003542 10.1371/journal.pgen.1003542 (2013).

44. Carrozza, M. J. et al. Histone H3 methylation by Set2 directs deacetylation of coding regions by Rpd3S to suppress spurious intragenic transcription. Cell 123, 581–592, doi:10.1016/j.cell.2005.10.023 (2005).

45. Joshi, A. A. & Struhl, K. Eaf3 chromodomain interaction with methylated H3-K36 links histone deacetylation to Pol II elongation. Mol Cell 20, 971–978, doi:10.1016/j.molcel.2005.11.021 (2005).

46. Luco, R. F. et al. Regulation of alternative splicing by histone modifications. Science 327, 996–1000, doi:10.1126/science.1184208 (2010).

47. Schwartz, S., Meshorer, E. & Ast, G. Chromatin organization marks exon-intron structure. Nat Struct Mol Biol 16, 990–995, doi:10.1038/nsmb.1659 (2009).

48. Zhou, H. L., Luo, G., Wise, J. A. & Lou, H. Regulation of alternative splicing by local histone modifications: potential roles for RNA-guided mechanisms. Nucleic Acids Res 42, 701–713, doi:10.1093/nar/gkt875 (2014).

49. Cattanach, B. M., Beechey, C. V. & Peters, J. Interactions between imprinting effects: summary and review. Cytogenet Genome Res 113, 17–23, doi:10.1159/000090810 (2006).

50. Fergusonsmith, A. C., Cattanach, B. M., Barton, S. C., Beechey, C. V. & Surani, M. A. Embryological and Molecular Investigations of Parental Imprinting on Mouse Chromosome-7. Nature 351, 667–670, doi:DOI 10.1038/351667a0 (1991).

51. Strogantsev, R. et al. Allele-specific binding of ZFP57 in the epigenetic regulation of imprinted and non-imprinted monoallelic expression. Genome Biol 16, doi:ARTN 112 10.1186/s13059-015-0672-7 (2015).

52. Lawson, H. A., Cheverud, J. M. & Wolf, J. B. Genomic imprinting and parent-of-origin effects on complex traits. Nat Rev Genet 14, 608–617, doi:10.1038/nrg3543 (2013).

53. Babak, T. et al. Genetic conflict reflected in tissue-specific maps of genomic imprinting in human and mouse. Nat Genet 47, 544–U158, doi:10.1038/ng.3274 (2015).

54. Baran, Y. et al. The landscape of genomic imprinting across diverse adult human tissues. Genome Res 25, 927–936, doi:10.1101/gr.192278.115 (2015).

55. Rivera-Mulia, J. C. et al. Allele-specific control of replication timing and genome organization during development. Genome Res 28, 800–811, doi:10.1101/gr.232561.117 (2018).

56. Maas, A. H. & Appelman, Y. E. Gender differences in coronary heart disease. Neth Heart J 18, 598–602, doi:10.1007/s12471-010-0841-y (2010).

57. Mosca, L., Barrett-Connor, E. & Wenger, N. K. Sex/gender differences in cardiovascular disease prevention: what a difference a decade makes. Circulation 124, 2145–2154, doi:10.1161/CIRCULATIONAHA.110.968792 (2011).

58. Group, E. U. C. C. S. et al. Gender in cardiovascular diseases: impact on clinical manifestations, management, and outcomes. Eur Heart J 37, 24–34, doi:10.1093/eurheartj/ehv598 (2016).

59. Hershberger, R. E., Morales, A. & Siegfried, J. D. Clinical and genetic issues in dilated cardiomyopathy: a review for genetics professionals. Genet Med 12, 655–667, doi:10.1097/GIM.0b013e3181f2481f (2010).

60. Gardeux, V., David, F. P. A., Shajkofci, A., Schwalie, P. C. & Deplancke, B. ASAP: a web-based platform for the analysis and interactive visualization of single-cell RNA-seq data. Bioinformatics 33, 3123–3125, doi:10.1093/bioinformatics/btx337 (2017).

61. Yousif, A., Drou, N., Rowe, J., Khalfan, M. & Gunsalus, K. C. NASQAR: a web-based platform for high-throughput sequencing data analysis and visualization. Bmc Bioinformatics 21, doi:ARTN 267 10.1186/s12859-020-03577-4 (2020).

62. de Soysa, T. Y. et al. Single-cell analysis of cardiogenesis reveals basis for organ-level developmental defects. Nature 572, 120–124, doi:10.1038/s41586-019-1414-x (2019).

63. Goodyer, W. R. et al. Transcriptomic Profiling of the Developing Cardiac Conduction System at Single-Cell Resolution. Circ Res 125, 379–397, doi:10.1161/CIRCRESAHA.118.314578 (2019).

64. Jia, G. et al. Single cell RNA-seq and ATAC-seq analysis of cardiac progenitor cell transition states and lineage settlement. Nat Commun 9, 4877, doi:10.1038/s41467-018-07307-6 (2018).

65. See, K. et al. Single cardiomyocyte nuclear transcriptomes reveal a lincRNA-regulated de- differentiation and cell cycle stress-response in vivo. Nat Commun 8, 225, doi:10.1038/s41467-017-00319-8 (2017).

66. Li, H. et al. The Sequence Alignment/Map format and SAMtools. Bioinformatics 25, 2078–2079, doi:10.1093/bioinformatics/btp352 (2009).

67. Heberle, H., Meirelles, G. V., da Silva, F. R., Telles, G. P. & Minghim, R. InteractiVenn: a web-based tool for the analysis of sets through Venn diagrams. Bmc Bioinformatics 16, 169, doi:10.1186/s12859-015-0611-3 (2015).

68. Khan, A. & Mathelier, A. Intervene: a tool for intersection and visualization of multiple gene or genomic region sets. Bmc Bioinformatics 18, 287, doi:10.1186/s12859-017-1708-7 (2017).

69. Raudvere, U. et al. g:Profiler: a web server for functional enrichment analysis and conversions of gene lists (2019 update). Nucleic Acids Res 47, W191–W198, doi:10.1093/nar/gkz369 (2019).

70. Chen, E. Y. et al. Enrichr: interactive and collaborative HTML5 gene list enrichment analysis tool. Bmc Bioinformatics 14, 128, doi:10.1186/1471-2105-14-128 (2013).

71. Kuleshov, M. V. et al. Enrichr: a comprehensive gene set enrichment analysis web server 2016 update. Nucleic Acids Res 44, W90–97, doi:10.1093/nar/gkw377 (2016).

72. Meers, M. P., Tenenbaum, D. & Henikoff, S. Peak calling by Sparse Enrichment Analysis for CUT&RUN chromatin profiling. Epigenet Chromatin 12, doi:ARTN 42 10.1186/s13072-019-0287-4 (2019).

